# Cell-type-specific co-expression inference from single cell RNA-sequencing data

**DOI:** 10.1101/2022.12.13.520181

**Authors:** Chang Su, Zichun Xu, Xinning Shan, Biao Cai, Hongyu Zhao, Jingfei Zhang

**Affiliations:** Department of Biostatistics, Yale University; Information Systems and Operations Management, Emory University

**Keywords:** cell-type-specific analysis, expression-measurement model, gene co-expression network, sequencing depth, single cell RNA-seq

## Abstract

The inference of gene co-expressions from microarray and RNA-sequencing data has led to rich insights on biological processes and disease mechanisms. However, the bulk samples analyzed in most studies are a mixture of different cell types. As a result, the inferred co-expressions are confounded by varying cell type compositions across samples and only offer an aggregated view of gene regulations that may be distinct across different cell types. The advancement of single cell RNA-sequencing (scRNA-seq) technology has enabled the direct inference of co-expressions in specific cell types, facilitating our understanding of cell-type-specific biological functions. However, the high sequencing depth variations and measurement errors in scRNA-seq data present significant challenges in inferring cell-type-specific gene co-expressions, and these issues have not been adequately addressed in the existing methods. We propose a statistical approach, CS-CORE, for estimating and testing cell-type-specific co-expressions, built on a general expression-measurement model that explicitly accounts for sequencing depth variations and measurement errors in the observed single cell data. Systematic evaluations show that most existing methods suffer from inflated false positives and biased co-expression estimates and clustering analysis, whereas CS-CORE has appropriate false positive control, unbiased co-expression estimates, good statistical power and satisfactory performance in downstream co-expression analysis. When applied to analyze scRNA-seq data from postmortem brain samples from Alzheimer’s disease patients and controls and blood samples from COVID-19 patients and controls, CS-CORE identified cell-type-specific co-expressions and differential co-expressions that were more reproducible and/or more enriched for relevant biological pathways than those inferred from other methods.

## 1 Introduction

The past two decades have seen great advances in gene co-expression studies using microarrays and RNA sequencing technologies, leading to rich insights on biological processes and disease mechanisms (Zhang and Horvath, 2005; Mostafavi et al., 2018; Koplev et al., 2022). To date, most co-expression analyses have been performed on bulk samples that are a mixture of different cell types. As a result, the inferred networks are confounded with varying cell type compositions across samples and limited to an aggregated view of gene regulations that may differ considerably across cell types (Heintzman et al., 2009; Su et al., 2022). To infer cell-type-specific networks from bulk samples, cell sorting can be performed, but the techniques are tedious and subject to technical artifacts (Box et al., 2020).

With scRNA-seq technology such as droplet-based methods, gene expressions can now be measured in individual cells with annotated cell types (Hao et al., 2021), offering a great opportunity to construct cell-type-specific co-expression networks. However, such an analytical task is challenged by the unique characteristics of scRNA-seq data such as their high sequencing depth variations and measurement errors. For scRNA-seq data, the expression level of a specific gene is measured through the observed UMI (unique molecular identifier) count for this gene, and the sequencing depth of a cell is the sum of UMI counts across all genes. For a typical single cell experiment, there is substantial variation of sequencing depths across cells (e.g., 400 - 20,000) (Hafemeister and Satija, 2019; Sarkar and Stephens, 2021). As a result, gene co-expressions measured via correlations of UMI counts across cells can be seriously confounded by varying sequencing depths, resulting in inflated false positive findings in detecting co-expressed gene pairs. This confounding issue cannot be addressed using standard normalization strategies, as will be shown later. Besides varying sequencing depths, measurement errors in the UMI count data pose an additional challenge in inferring co-expression levels as the errors tend to attenuate correlation estimates with different degrees for genes with different expression levels.

Recent years have seen the developments of a number of methods to better capture coexpressions from scRNA-seq data than a simple normalization-based approach, including locCSN (Wang et al., 2021b), Noise Regularization (Zhang et al., 2021), Normalisr (Wang, 2021), propr (Quinn et al., 2017), and SpQN (Wang et al., 2022). These methods consider novel association metrics or additional adjustments when inferring co-expressions from scRNA-seq data. However, the proposed procedures do not have rigorous justifications as they are not explicitly based on the underlying data generating mechanisms and do not appropriately account for measurement errors and varying sequencing depths across cells. Besides the above co-expression estimation methods, a recently proposed method called sctransform (Hafemeister and Satija, 2019) estimates gene expression levels from scRNA-seq data by removing the effect of varying sequencing depths via negative binomial regressions. Although sctransform was not developed for co-expression estimation, one sensible approach is to calculate correlations of expression levels that have been adjusted for sequencing depths by sctransform; we refer to this approach as *ρ*-sctransform in our following discussion. As will be demonstrated later, the sequencing depth normalization in sctransform, designed to infer expression levels, can be inadequate in removing biases from sequencing variation and measurement errors when inferring co-expressions. In our systematic evaluations of different methods based on simulated and permuted real scRNA-seq data, we found that all the other methods, including *ρ*-sctransform, suffer from inflated type-I errors, varying degrees of estimation biases, reduced accuracy in detecting co-expressions, and potentially misleading results in downstream co-expression analysis such as clustering and principal component analysis.

Here, we present a statistical approach for estimating and testing co-expressions from scRNA-seq data, called CS-CORE (cell-type-specific co-expressions). Specifically, CS-CORE models the unobserved true gene expression levels as latent variables, linked to the observed UMI counts through a measurement model that accounts for both sequencing depth variations and measurement errors. Under this model, CS-CORE implements a fast and efficient iteratively re-weighted least squares approach for estimating the true correlations between underlying expression levels, together with a theoretically justified statistical test to assess whether two genes are independent. The proposed model in CS-CORE does not impose any distributional assumptions on the underlying expression levels and can flexibly accommodate single cell data generating mechanisms such as negative binomial distributed counts. Through systematic evaluations based on simulated and permuted real scRNA-seq data, we found that CS-CORE had proper type-I error control, unbiased co-expression estimates and increased statistical power compared with other methods. CS-CORE also had satisfactory performance in downstream co-expression analysis.

We evaluated the utility of CS-CORE by applying it to multiple scRNA-seq data sets including postmortem brain samples from Alzheimer’s disease patients and controls (Lau et al., 2020) and peripheral blood mononuclear cells (PBMC) of COVID-19 patients and controls (Wilk et al., 2020). For both diseases, CS-CORE identified co-expressions that were more reproducible across independent data sets and more enriched with known transcription factor-target gene pairs than other methods. Clustering analysis using results from CS-CORE extracted co-expressed and differentially co-expressed gene modules that were more strongly enriched for relevant cell-type-specific biological functions than those inferred from other methods, highlighting the potential of CS-CORE in characterizing cell-type-specific biological functions and uncovering novel disease-related cell-type-specific pathways.

## 2 Results

### 2.1 Overview of CS-CORE

We have *n* cells with the observation for cell *i*, *i* =1, …, *n*, denoted by a vector (*x*_*i*1_, …, *x_ip_*) corresponding to the observed UMI counts for *p* genes. We use 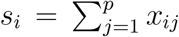 to denote the sequencing depth of cell *i*, which is the sum of UMI counts across all genes in this cell. Let (*z*_*i*1_, …, *z_ip_*) denote the underlying expression levels from *p* genes in cell *i*, defined to be the number of molecules from each gene relative to the total number of molecules in a cell (Sarkar and Stephens, 2021). Assume that

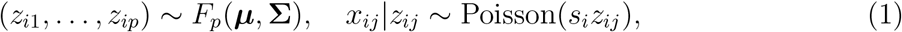

where *F_p_*(***μ***, **Σ**) is an unknown nonnegative *p*-variate distribution with mean vector ***μ*** = (*μ*_1_, …, *μ_p_*), 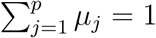, and covariance matrix 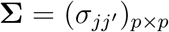. Here, *x_ij_* is the UMI count of gene *j* in cell *i*, assumed to follow a Poisson measurement model (Sarkar and Stephens, 2021) depending on the underlying expression level *z_ij_* and sequencing depth *s_i_*. This Poisson measurement model explicitly accounts for the sequencing depths and measurement errors. While a marginal expression-measurement model has been considered for modeling expression levels in bulk RNA-seq (Robinson et al., 2010; Love et al., 2014) and scRNA-seq data (Wang et al., 2018; Hafemeister and Satija, 2019; Townes et al., 2019), a joint expression-measurement model such as (1) is needed to infer co-expressions. Under (1), if *z_ij_* follows a Gamma distribution, then *x_ij_* follows a negative binomial distribution marginally.

We measure gene co-expressions by **Σ**_*p*×*p*_, which quantifies the correlation strength between the underlying expression levels. This definition of co-expression is precise and not biased by sequencing depth variations and measurement errors. Specifically, for any gene pair (*j, j*′), we measure co-expression via their correlation 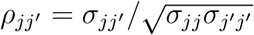.

Given UMI counts 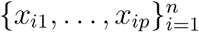 and sequencing depths 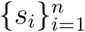, estimating the covariance matrix **Σ**_*p*×*p*_ is a challenging task. Without placing distributional assumptions on *F_p_*, we propose a moment-based iteratively reweighted least squares (IRLS) estimation procedure, that is fast to implement and statistically efficient. For each pair (*j, j*′), we also develop a theoretically justified hypothesis testing procedure that evaluates the independence between their expression levels *z*_*ij*_ and *z_ij′_*. The test statistic can be easily computed using IRLS estimates, does not require any distributional assumptions on *F_p_*, and follows a standard normal distribution under the null.

Details of the above estimation and testing procedures are given in Section 4.1. In summary, CS-CORE takes UMI counts and sequencing depths across cells as input and estimates correlations of the underlying expression levels as well as *p*-values for testing independence between gene pairs, without needing parameter tuning. The procedure removes the confounding effects of varying sequencing depths and the bias from measurement errors when inferring co-expressions, is theoretically justified and fast to implement.

### 2.2 CS-CORE has better control of false positive rates

To evaluate the performance of CS-CORE and illustrate the confounding effects from sequencing depth variations on other methods for independent gene pairs, we generated null data sets, where genes are not expected to co-express, by permuting the single nucleus RNA-seq (snRNA-seq) data from Lau et al. (2020) while making the sequencing depths across cells either constant or varying. Specifically, we normalized gene expressions (UMI counts) within each cell by its sequencing depth and, for each gene, we randomly permuted its normalized expression levels across cells. Then, we obtained UMI counts for each gene based on prespecified sequencing depth of each cell (Section 4.3). To examine effects of sequencing depth variations, we considered two settings with one set to observed sequencing depths in real data, which are highly variable, and one set to be constant across cells. This permutation procedure de-correlated gene expressions such that the average co-expression for each gene in the permuted data, calculated by averaging its co-expressions with all other genes, is expected to center around zero, regardless of sequencing depth variations.

We compared CS-CORE to other approaches, including locCSN (Wang et al., 2021b), Noise Regularization (Zhang et al., 2021), Normalisr (Wang, 2021), Pearson correlation, propr (Quinn et al., 2017), *ρ*-sctransform (Hafemeister and Satija, 2019), Spearman correlation and SpQN (Wang et al., 2022) (Section 4.2). Among these approaches, statistical tests for co-expressions are possible for Noise Regularization, Normalisr, Pearson correlation, *ρ*-sctransform and Spearman correlation.

For null data with high variations in sequencing depths, we found that co-expression estimates from most methods were biased with estimated average gene co-expressions different from zero (Figure 1A). The amount of bias varied with the expression level with distinct patterns for different methods. Meanwhile, in null data with no sequencing depth variations, there were minimal biases for these methods (Figure 1A), demonstrating that co-expression estimates can be biased by sequencing depth variations. By contrast, average co-expressions estimated by CS-CORE were unbiased and centered around zero, regardless of sequencing depth variations (Figure 1A). We observed the same qualitative patterns in our experiments with simulated data (Figure S2). One main cause of bias from other methods is no or inadequate adjustments of sequencing depth variations when measuring co-expressions, including the standard log transformations (e.g., Pearson, locCSN) and post-hoc adjustments (e.g., SpQN). We further illustrate this with additional simulations in Figure S1.

**Figure 1:**
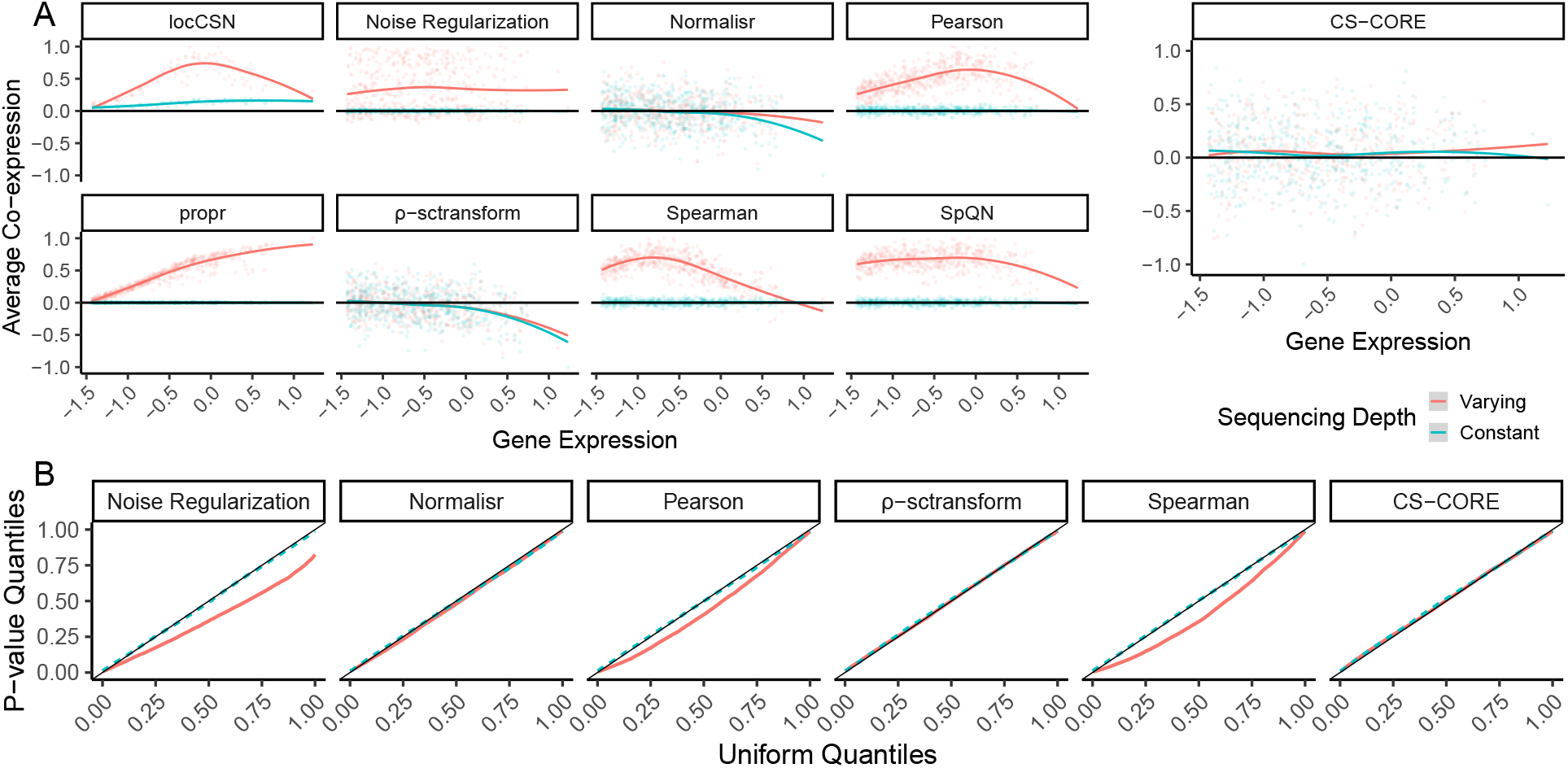
Validation of CS-CORE using permuted snRNA-seq data from Lau et al. (2020). Results from permuted data with varying and constant sequencing depths are colored with light red and blue, respectively. (A) Scatter plots with fitted curves showing expression (x-axis) and average co-expression (y-axis) of each gene with co-expression estimated using locCSN, Noise Regularization, Normalisr, Pearson correlation, propr, *ρ*-sctransform, Spearman correlation, SpQN and CS-CORE. Average co-expressions are re-scaled by the maximum value to aid comparison. (B) Q-Q plots comparing *p*-values for testing co-expressions of gene pairs against Uniform(0,1) using six methods with statistical tests, including Noise Regularization, Normalisr, Pearson correlation, *ρ*-sctransform, Spearman correlation and CS-CORE.

We also considered statistical tests for co-expressions in the permuted data. As the null hypothesis of no co-expression is expected to hold after permutation, *p*-values for testing independence of gene pairs should follow the Uniform[0,1] distribution. We found that in null data with no variations in sequencing depths, most methods had well-controlled type-I errors as the Q-Q plots showed matching quantiles between empirical distributions of *p*-values and the uniform distribution (Figure 1B). However, in null data with high variations in sequencing depths, Noise Regularization, Pearson and Spearman had inflated type I errors, demonstrating the confounding effects of sequencing depth variations (Figure 1B). While Normalisr and *ρ*-sctransform had controlled type-I errors with sequencing depth variations, they had biases in estimating co-expressions (Figure 2A). We also found that these two methods had reduced power in detecting co-expressions when compared to CS-CORE (Appendix, Figure S3).

**Figure 2:**
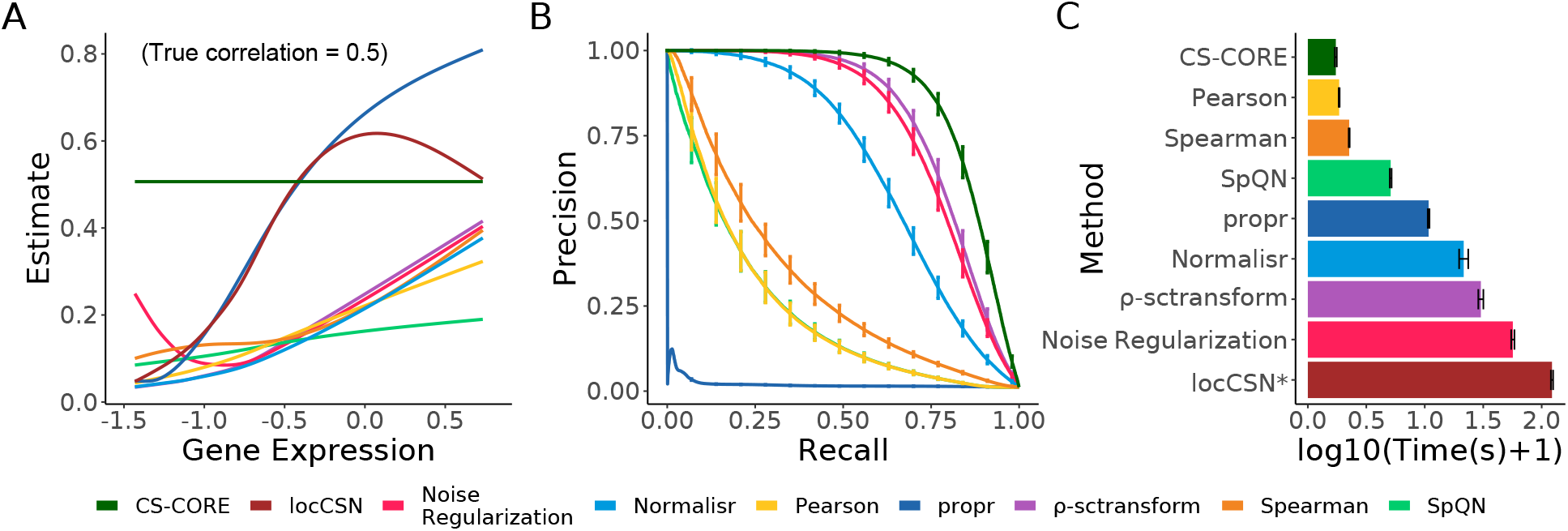
Validation of CS-CORE using simulated data, compared to locSCN, Noise Regularization, Normalisr, Pearson correlation, propr, *ρ*-sctransform, Spearman correlation and SpQN. (A) Curve-fitted co-expression estimates against geometric mean expression levels on gene pairs simulated with a true correlation of 0.5 (5,000 genes and 1,000 cells). (B) Precision-recall curves evaluated using 500 genes, 5,000 cells and a sparse co-expression matrix estimated from real data. Cut-off values are based on *p*-values for CS-CORE, Noise Regularization, Normalisr, Pearson correlation, *ρ*-sctransform, Spearman correlation and absolute values of co-expression estimates for propr and SpQN, as they are not equipped with statistical tests; locCSN is excluded due to its extreme demand in computing time. (C) Running times evaluated under the same setting as in (B). locCSN is evaluated using 0.2% of the cells to reduce computing time. The simulations were run on an Intel Xeon Gold 6240 @ 2.60GHz with one node and 50GB memory. The error bars denote one standard deviation across 100 replications.

### 2.3 CS-CORE has better co-expression estimation and detection accuracy

We evaluated the accuracy of CS-CORE in estimating and detecting co-expressions and illustrated another issue often referred to as the mean-correlation bias (Crow et al., 2016; Wang et al., 2022) in co-expression estimation. The mean-correlation bias is a separate issue from the confounding effect of varying sequencing depths. It arises, as measuring associations of the observed UMI counts, which profile the underlying expressions with measurement errors, tend to yield attenuated estimates due to the added errors. The amount of attenuation bias tends to decrease as the expression level increases (see Section 4.4) and correlations tend to be more accurately estimated for highly expressed genes. As a result, highly expressed genes can appear more correlated as an artifact. This attenuation bias has also been noted in analyzing bulk RNA-seq data (Saccenti et al., 2020; Wang et al., 2022), but it can be exacerbated by the shallow sequencing depths frequently seen in scRNA-seq data.

To demonstrate this, we simulated expression data for gene pairs with varying expression levels and a correlation of *ρ* = 0.5 following marginal negative binomial distributions (see Section 4.5). For co-expressed gene pairs with a true correlation of 0.5, we found that correlation estimates from all other methods were inaccurate (Figure 2A) with most methods severely underestimating co-expressions for genes with low or moderate expression levels. The correlation estimates also spuriously increased with expression levels for most methods. By contrast, CS-CORE could accurately estimate co-expressions (Figure 2A) and was not subject to mean-correlation bias. This is because CS-CORE is based on an expression-measurement model and explicitly measures co-expressions using correlations of the underlying expression levels, free of measurement errors. The mean-correlation bias remained on data simulated with no variations in sequencing depths (Figure S4), suggesting that the mean-correlation bias is a separate source of bias from varying sequencing depths. We further evaluated the co-expression detection accuracy in simulations with *p* = 500 where co-expressed pairs were set to those inferred from real data (see Section 4.5). The precision-recall curves in Figure 2B show that CS-CORE achieves the highest area-under-the-curve value.

Finally, we compared the computing time of different methods (Figure 2C) under the simulation setting considered in Figure 2B. It is seen that CS-CORE is highly computationally efficient as it uses a least squares estimation procedure. Specifically, CS-CORE was faster to implement than the state-of-the-art method, locCSN, which is based a local nonparametric test and *ρ*-sctransform, which requires fitting marginal negative binomial regressions using likelihood-based approaches. The computing time of CS-CORE is comparable to simple procedures such as Pearson and Spearman, as they both include a normalization step (see Section 4.2).

### 2.4 Other methods can lead to biased results in downstream co-expression analysis

Bias in estimating co-expressions can negatively impact important downstream co-expression analyses such as clustering and principal component analysis (PCA). To evaluate the performance of CS-CORE and other methods on such downstream analytical tasks, we simulated *n* = 2, 000 cells for *p* = 100 genes with varying expression levels and a co-expression matrix with four clusters (see Section 4.5 and S2). We estimated co-expression networks using CS-CORE and other methods, and compared them to the true co-expression network (Figure 3A). In particular, when plotting the results from each method, we ordered the genes by applying hierarchical clustering to the estimated co-expression network. We found that CS-CORE was the only method that could accurately estimate co-expressions and be used to recover true clusters. The estimated co-expression networks and inferred cluster labels from other methods were strikingly inaccurate. These findings were further supported by evaluating the clustering accuracy (Figure 3B), measured using adjusted Rand index, and the accuracy in estimating the top principal components (Figure 3C), measured using subspace distance (Golub and Van Loan, 2013).

**Figure 3:**
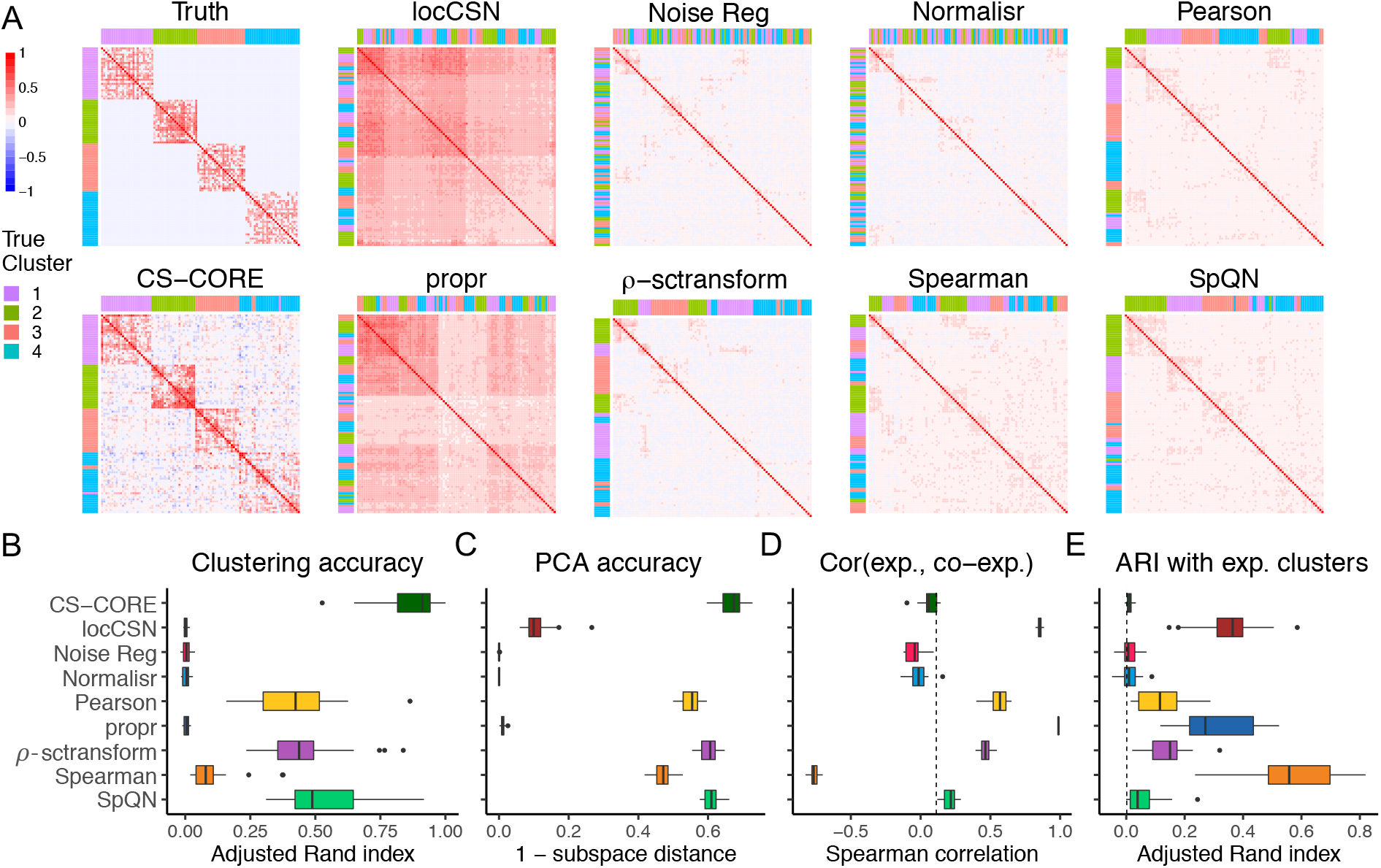
Evaluation of CS-CORE in recovering clusters and principal components using simulated data, compared to locCSN, Noise Regularization, Normalisr, Pearson correlation, propr, *ρ*-sctransform, Spearman correlation and SpQN. (A) Heatmaps of true and estimated co-expression networks from simulations. When plotting results from each method, genes were ordered by applying hierarchical clustering to the estimated co-expression network and color coded by their true cluster labels. (B) Adjusted Rand index (ARI) between true clusters and clusters extracted from co-expression networks estimated using different methods. (C) Accuracy in recovering principal components, calculated using subspace distance (Golub and Van Loan, 2013) between the top four singular vectors of the true co-expression matrix and those of the estimated co-expression matrix. (D) Spearman correlations between the expression levels and estimated average co-expression levels of genes, with ground truth calculated from simulation settings marked with a dashed line. (E) ARI between clusters extracted from estimated co-expression networks and clusters extracted from clustering gene expression levels, with the true ARI calculated from parameters used in simulation settings marked with a dashed line. (B)-(E) were evaluated with 25 replications.

To highlight the mean-correlation bias, we computed the correlation between gene expression levels and estimated co-expression levels. As expression levels were randomly assigned independent of correlation strengths, the true correlation between gene expression and coexpression levels should be close to zero, as marked in Figure 3D. However, we found that the co-expression levels estimated from locCSN, Pearson correlation, propr, *ρ*-sctransform and Spearman correlation were spuriously correlated with the mean expression level. One implication of this mean-correlation bias is that, as highly expressed genes often appear highly co-expressed with other genes as an artifact, clustering methods tend to incorrectly cluster genes with similar expression levels in a co-expression cluster and expression levels become falsely predictive of the network modules (Figure 3E). In another data example, we demonstrated that this mean-correlation bias could also lead to spurious clustering structures on null data where genes are not co-expressed (Figure S5).

### 2.5 CS-CORE identified more reproducible and biologically relevant co-expressions from AD and control brain samples

We applied CS-CORE to a snRNA-seq data set collected from the prefrontal cortical regions of 12 Alzheimer’s disease (AD) patients and nine controls in Lau et al. (2020). In particular, we focused our comparison with *ρ*-sctransform, as it gives the best overall performance in Figure 1–3 and allows for statistical tests.

First, using samples from controls, we estimated the co-expression network among top 5,000 highly expressed genes in five major brain cell types including astrocyte (Ast), excitatory neuron (Ex), inhibitory neuron (In), oligodendrocyte (Oli) and microglia (Mic), and evaluated the reproducibility of identified co-expressions using two independent snRNA-seq data sets on prefrontal cortex from Mathys et al. (2019) and Morabito et al. (2021) (Section S3.3). Figure 4A shows that the co-expressed gene pairs inferred by CS-CORE were more reproducible in Mathys et al. (2019) than those inferred by *ρ*-sctransform across different *p*-value cutoffs and cell types, suggesting CS-CORE has greater statistical power to detect true co-expression signals. We had similar observations for data from Morabito et al. (2021) (Figure S6).

**Figure 4:**
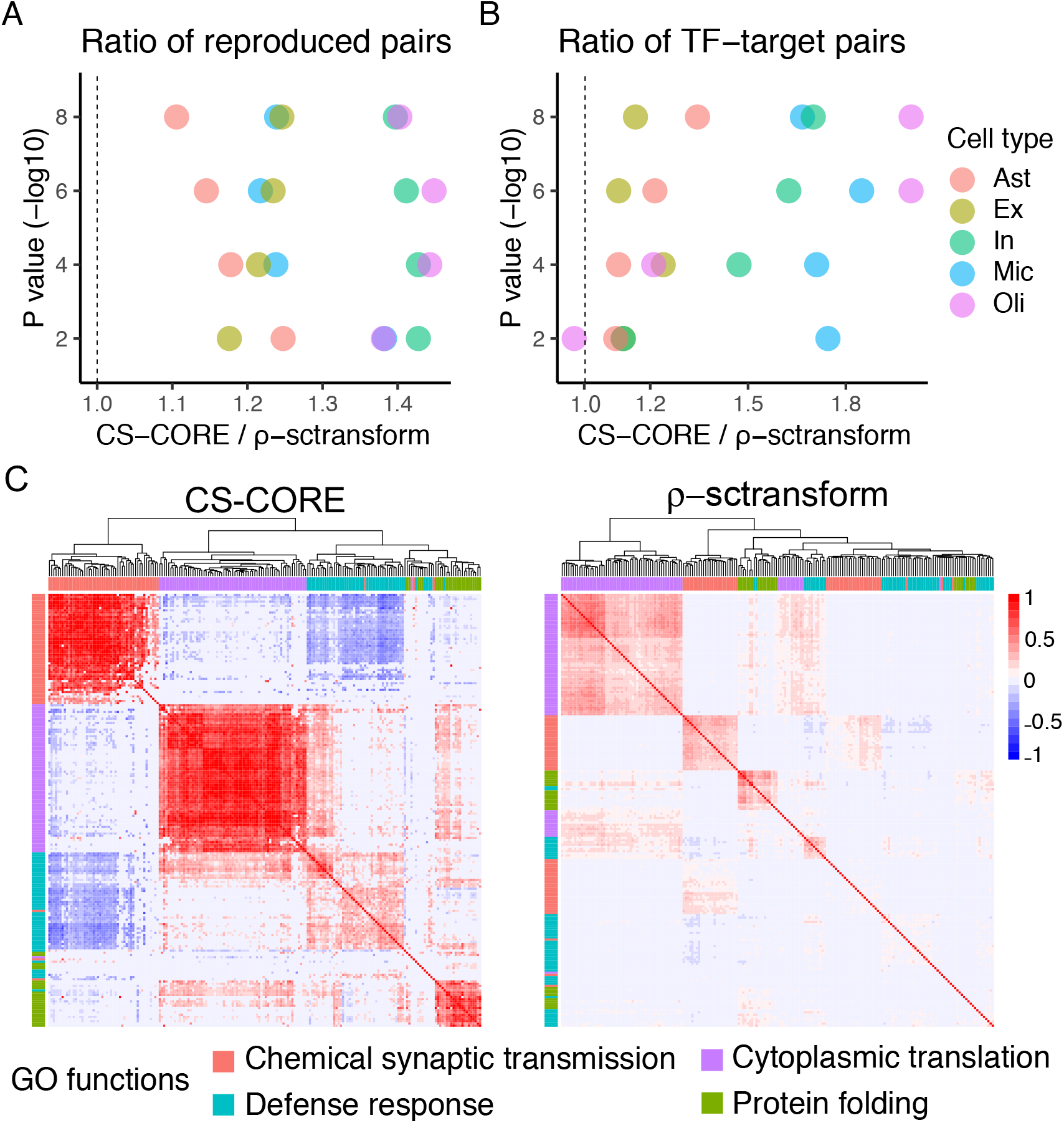
Co-expression analysis using AD brain samples in Lau et al. (2020). We used the cells in five major brain cell types from control subjects from Lau et al. (2020) to estimate cell-type-specific co-expression networks. **A.** Ratio of the numbers of gene pairs that were identified as significant in both Lau et al. (2020) and Mathys et al. (2019) at specified *p*-value cutoffs between CS-CORE and *ρ*-sctransform. **B.** Ratio of the numbers of gene pairs that were identified as significant and overlapped with known TF-target gene pairs in the TRRUST database (Han et al., 2018) between CS-CORE and *ρ*-sctransform. **C.** Heatmaps of microglia-specific co-expression network estimates on genes from four GO terms on microglia’s functions with genes ordered by hierarchical clustering.

Next, by evaluating the overlap of co-expressed pairs with a database on known Transcription Factor(TF)-target gene pairs (Han et al., 2018), we found CS-CORE recovered more known TF-target pairs than *ρ*-sctransform from the inferred networks (Figure 4B). Additionally, we extracted co-expressed gene modules by applying WGCNA (Langfelder and Horvath, 2008) on significantly co-expressed gene pairs, which were then evaluated using Gene Ontology (GO) enrichment analysis (Wu et al., 2021) (see Section S3.1). Our enrichment analysis used the 5,000 highly expressed genes as the background gene set, such that enrichment of any module is not attributed to its high expression levels. For microglia, the innate immune brain cells with a central role in the AD neuroinflammation mechanism (Heneka et al., 2015), clustering based on CS-CORE identified four modules strongly enriched for GO terms related to microglia’s functions, including defense response, chemical synaptic transmission, cytoplasmic translation and protein folding, respectively, while only two of these four functions were found enriched for modules inferred based on *ρ*-sctransform with less significant *p*-values and lower gene ratios (Tables S2, S3). In particular, Figure 4C shows the estimated co-expression networks, with genes ordered by hierarchical clustering, on a subset of genes from the four GO terms. It is seen that CS-CORE accurately grouped genes into respective biological functions, with genes in the same GO function densely connected. By contrast, *ρ*-sctransform only partially recovered some gene modules and the estimated co-expressions are generally much weaker. Besides microglia, CS-CORE also identified gene modules that were enriched for cell-type-specific functions in astrocytes (synaptic signalling, protein folding, cellular response to hypoxia), inhibitory neurons (synaptic membrane) and oligodendrocytes (cell-cell signaling, cholesterol metabolic process), while these functions were either not or much less enriched for modules inferred based on *ρ*-sctransform (Table S2, S3). This further highlights the potential of CS-CORE in uncovering cell-type-specific biological pathways.

Finally, we constructed the differential co-expression network in microglia between AD patients and controls from Lau et al. (2020) to investigate the biological pathways dysregulated in AD (see Section 4.6). We applied clustering analysis to the differential network to extract gene modules that shared similar co-expression changes in AD and performed GO enrichment analysis. Clustering based on CS-CORE identified three differentially coexpressed gene modules enriched for cell-type-specific functional pathways that are implied in AD disease mechanisms, including protein folding (Roychaudhuri et al., 2009), synapse signaling transduction (Kamat et al., 2016), and protein kinase (toll-like receptors) signaling pathways (Landreth and Reed-Geaghan, 2009) (Table S4). In comparison, *ρ*-sctransform did not identify any differentially co-expressed module enriched with cell-type-specific biological or disease-related functions (Table S5).

### 2.6 CS-CORE identified upregulated co-expressions in Interferon signalling pathway from COVID-19 blood samples

We applied CS-CORE to a scRNA-seq data set from human peripheral blood mononuclear cells (PBMC) of seven hospitalized patients with SARS-CoV-2 and six controls (Wilk et al., 2020) to identify biological pathways differentially regulated in COVID-19 patients.

Using samples from controls, we estimated cell-type-specific co-expressions among the top 5,000 highly expressed genes in five major immune cell types, including B cells, CD4 positive T cells, CD8 positive T cells, monocytes and natural killer (NK) cells. Using an independent scRNA-seq data set on PBMC (Unterman et al., 2022), we found that CS-CORE yielded a larger number of reproducible co-expressed gene pairs than *ρ*-sctransform across different *p*-value cutoffs and cell types (Figure S7A). CS-CORE also uncovered more gene pairs that overlapped with known TF-target gene pairs and more gene modules with stronger cell-type-specific functional enrichment than *ρ*-sctransform across cell types through GO enrichment analysis (Figure S7B, Tables S6-S10). For example, CS-CORE identified three co-expression modules enriched for the biological functions of B cells, including antigen processing via MHC Class II, adaptive immune response and response to inteferon-alpha (Table S6). In contrast, only one of these three functions was found enriched in a module inferred based on *ρ*-sctransform with a less significant *p*-value and a lower gene ratio (Table S9). Our results on PBMC again show that CS-CORE can recover biologically more meaningful co-expressions than other methods.

We next investigated cell-type-specific responses to SARS-CoV-2 viral infection in monocytes using a differential co-expression analysis similar to the one performed in the previous section between AD patients and controls. Clustering analysis revealed gene modules that share similar co-expression changes in monocytes in response to SARS-CoV-2. In particular, three gene modules inferred using co-expression estimates from CS-CORE were significantly enriched for immune responses based on GO enrichment analysis, including inflammatory response, virus defense response, and cellular stress response (Table S11). In contrast, *ρ*-sctransform only identified one gene module associated with virus defense response with much weaker enrichment signals (Table S12). In Figure 5, we highlight a module identified by CS-CORE, which is enriched for the interferon signalling pathway (Table S13), a key immune signature in COVID-19 patients that has been demonstrated in multiple studies (Acharya et al., 2020; Hadjadj et al., 2020; Lee and Shin, 2020). While it is known that the expression levels of interferon-stimulated genes are upregulated in monocytes from COVID-19 patients, by comparing the CS-CORE estimates in monocytes between COVID-19 patients and controls, we identified upregulated co-expressions among interferon-stimulated genes, suggesting increased gene coordination in the interferon signalling pathway upon viral infection. We also found stronger co-expressions between genes in the interferon signaling and antigen presentation pathways among COVID-19 patients, suggesting stronger concerted immune responses between these two pathways. Finally, we note that this gene module also contains multiple known genes in the SARS-CoV-2 infection Reactome pathway, revealing cell-type-specific changes in co-expressions among known disease-related genes.

**Figure 5:**
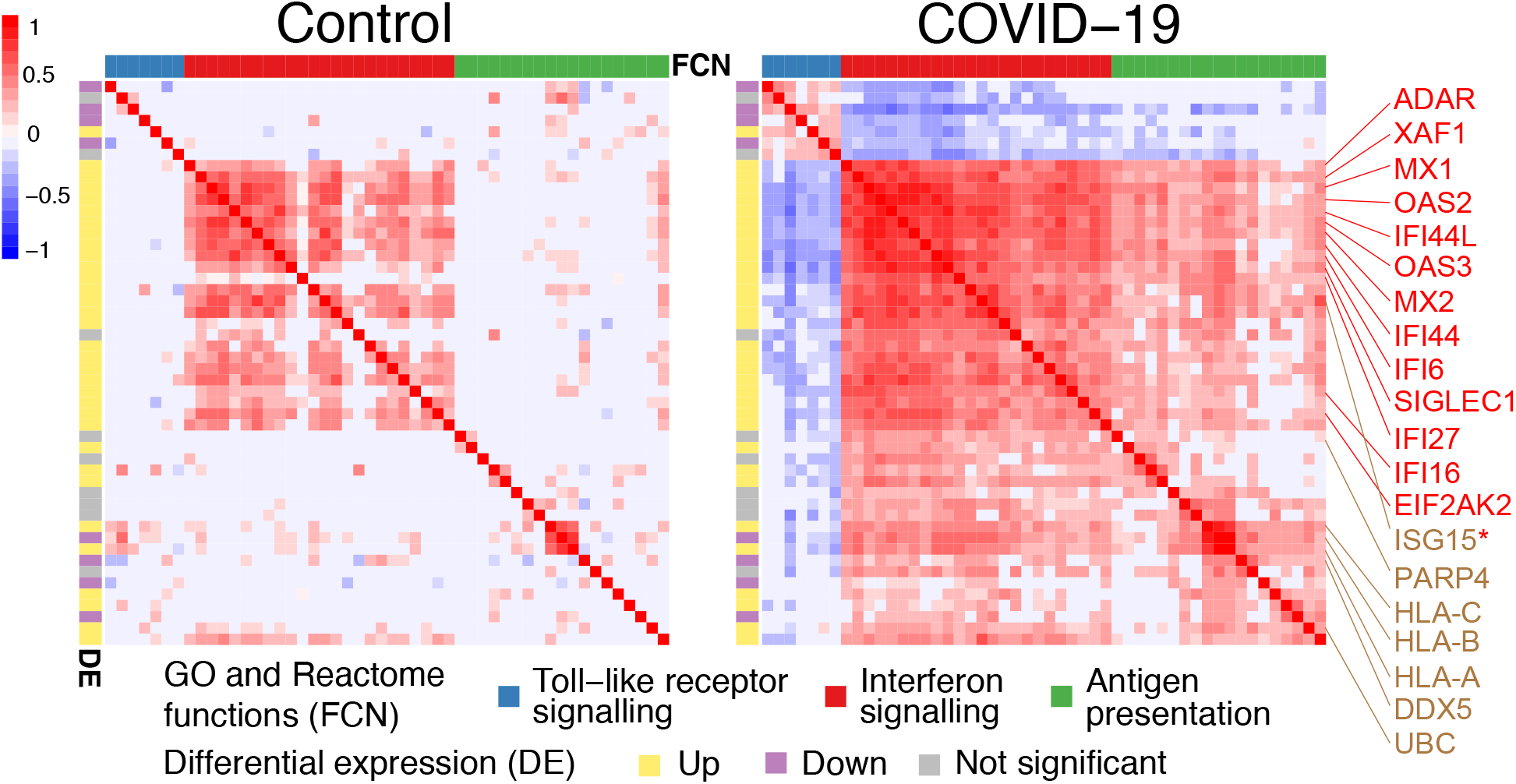
CS-CORE estimates in monocytes from control subjects and COVID-19 patients. Known interferon-stimulated genes are colored in red. Genes in the SARS-CoV-2 infection Reactome pathway are colored in brown. * is used to mark genes that belong to both gene sets. We performed a differential co-expression analysis on top 1,000 highly expressed genes in monocytes and obtained modules of genes that shared similar changes in co-expressions between cells from COVID-19 patients and controls. For a differentially co-expressed gene module enriched for the interferon signalling pathway, we focused on genes that had strong differential signals (sum of absolute differential co-expressions greater than the median) and visualized the co-expression network estimates in control subjects and COVID-19 patients.

## 3 Discussion

We developed a comprehensive statistical approach, CS-CORE, for estimating and testing cell-type-specific co-expressions based on scRNA-seq data. CS-CORE adopts a multivariate expression-measurement model for the observed UMI counts and a pair-wise IRLS method for estimation and testing. It does not place distributional assumptions on the underlying expression levels and can be implemented very efficiently to estimate and test co-expressions in a large network. We demonstrated the better performance of CS-CORE than other methods through both simulations and real data analyses.

Our work pointed to two potential sources of biases when inferring co-expressions from UMI counts. The first one is the varying sequencing depths across cells, which can lead to inflated false positive findings in detecting co-expressions, as a pair of independent genes may appear co-expressed as a result of the sequencing depth variations across cells. The second one is the error from the measurement process, causing the observed UMI counts to deviate from the underlying expression levels. Under the Poisson measurement model, this deviation is a function of both the expression level and the sequencing depth. When estimating the underlying co-expression level for a pair of genes, correlations between UMI counts tend to be biased towards zero as a result of the measurement errors. In our experiments, we observed such an attenuation bias in most methods we compared to, leading to inaccuracy and reduced power in estimating and detecting co-expressions. These two distinct sources of biases, when combined, cause serious issues in estimating and testing for co-expressions. As demonstrated in our analysis, no other methods can adequately address both. Our approach CS-CORE addresses them by explicitly modeling the measurement process, accounting for both varying sequencing depths and measurement errors, and estimates the first and second moments of the underlying multivariate expression model to produce estimates of co-expressions, without any specific distributional assumptions.

There has been recent work that makes cell-type-specific inferences from bulk samples leveraging cell type deconvolution techniques (Jin et al., 2021; Wang et al., 2021a). These work often aims to estimate cell-type-specific expressions and compositions in bulk samples (Wang et al., 2019; Newman et al., 2019; Jaakkola and Elo, 2021; Cai et al., 2022). In particular, a recent method CSNet (Su et al., 2022) focuses on estimating cell-type-specific co-expressions from bulk sample data. The rich bulk samples collected over past decades and the increasingly available scRNA-seq data together offer a great opportunity to integrate bulk samples and single cell data to draw cell-type-specific inferences of co-expressions. The proposed method CS-CORE provides a useful tool in developing methods for such integrative analyses.

In CS-CORE, we have assumed that gene expressions from cells of the same cell type follow the same distribution. This assumption may not hold when the cells are collected from individuals with different genetic, demographic and clinical characteristics. For example, there is a growing interest in studying the genetic basis of cell-type-specific gene expression and co-expression differences across individuals using single cell data, and such population level single cell data are becoming increasingly available (Young et al., 2021; Nathan et al., 2022). As an important next step, we plan to extend the CS-CORE framework to infer individualized cell-type-specific co-expression networks and to study the differences in gene co-expressions across genotypes and conditions, shedding light on individualized and context-specific biological functions and pathways.

In summary, the CS-CORE method introduced in this article is statistically sound and computationally efficient. Compared to the other methods, it generates more reproducible and biologically more relevant cell-type-specific co-expression networks across multiple scRNA-seq data sets. With the rapid increase of scRNA-seq studies, we believe that CS-CORE offers a powerful and robust statistical tool to infer cell-type-specific co-expression networks to characterize biological pathways and molecular mechanisms at the cell type level.

## 4 Methods

### 4.1 CS-CORE method

Under the expression-measurement model defined in (1), it holds that 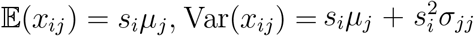, and 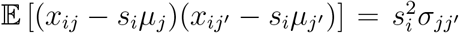. This motivates us to estimate *μ_j_*’s and 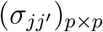 via the following set of regression equations:

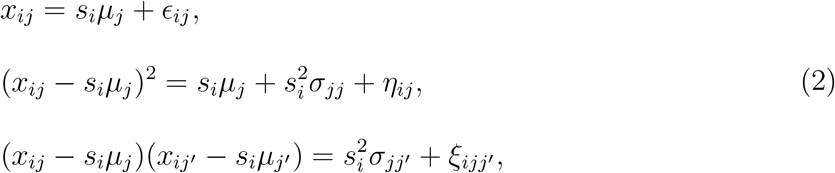

where *ϵ_ij_*, *η_ij_*, and *ξ_ijj′_* are independent and mean-zero error variables for all *i*, *j*, *j*′. Specifically, given UMI counts *x_ij_*’s and sequencing depths *s_i_*’s, the mean parameter *μ_j_* is estimated via 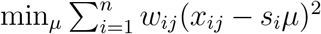, where *w_ij_* is the weight for cell *i* to be determined. Given the estimates 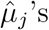, we estimate *σ_jj_* and 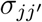 with 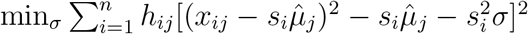 and 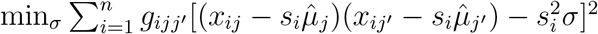, respectively, where *h_ij_* and *g_ijj′_* are weights to be determined. These weighted least squares can be computed very efficiently.

In CS-CORE, we carefully select and update the weights via an IRLS procedure, such that the weighted least squares estimators are statistically efficient; see the detailed procedure in Algorithm S1. The most ideal weights, in terms of statistical efficiency, should be the reciprocal of the variances of the error variables in (2). Hence, we set 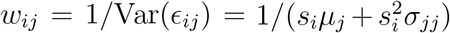, which is updated in each step of the IRLS estimation. The analytical forms of Var(*η_ij_*) and Var(*ξ_ijj′_*) are difficult to derive as we do not place distributional assumptions on *z_ij_*. Given weights *w_ij_*’s for the mean parameter estimation, we set weights for variance and covariance estimation as 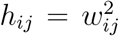 and 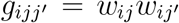, respectively, which yield good performance in our experiments and the IRLS procedure typically converges within five iterations. In practice, we add a regularization step to the variance parameters *σ_jj_*’s used in calculating the weights, as their estimates can be variable, leading to highly variable weights.

Next, we develop a statistical test to assess whether a gene pair have independent expression levels. Under model (1) and when *z_ij_* and *z_ij′_* are independent, 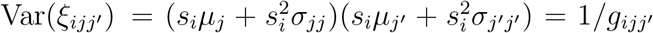. Letting 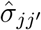 be estimated with true *μ_j_*’s, we define the test statistic 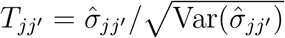, which can be calculated as

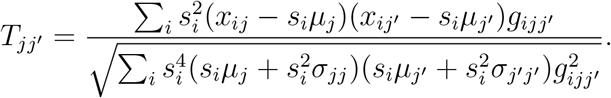

It then follows that *T_jj′_* ~ *N*(0,1) under the null hypothesis that *z_ij_* and *z_ij′_* are independent. This result allows us to directly compute *p*-values by plugging in IRLS estimated *μ_j_*’s and *σ_jj_*’s, all of which are consistent weighted least squares estimators.

### 4.2 Other co-expression estimation and testing methods using scRNA-seq data

We compared CS-CORE with eight other methods for inferring gene co-expression from single cell data, including locCSN (Wang et al., 2021b), Noise Regularization (Zhang et al., 2021), Normalisr (Wang, 2021), Pearson correlation, propr (Quinn et al., 2017), *ρ*-sctransform, Spearman correlation and SpQN (Wang et al., 2022). The method locCSN was applied on log normalized data log(*x_ij_/s_i_* + 1) and computed with the Python implementation provided at https://github.com/xuranw/locCSN. While locCSN estimates one network per cell, we followed the authors’ instructions to aggregate cell-specific co-expressions into celltype-specific co-expressions, as stated in Wang et al. (Wang et al., 2021b) that averaging provides stable estimates of the network structure. The method propr refers to *ρ_p_* in Quinn et al. (2017) and was calculated with the R package ‘propR’ (v.4.2.6). For *ρ*-sctransform, we computed the residuals of sctransform using R package Seurat (v.4.0.3) and evaluated Pearson correlations between the residuals. The Spearman (Pearson) correlation was calculated on log normalized expression data using the R package ‘stats’ (v.4.1.3). Noise Regularization (Zhang et al., 2021) was implemented from https://github.com/RuoyuZhang/ NoiseRegularization, Normalisr (Wang, 2021) was computed with the Python implementation from https://github.com/lingfeiwang/normalisr (v.1.0.0) and SpQN (Wang et al., 2022) was computed with R package ‘SpQN’ (v.1.6.0).

Among the above eight methods, statistical tests for co-expressions are possible for Noise Regularization, Normalisr, Pearson correlation, *ρ*-sctransform and Spearman correlation. Specifically, test statistics for Noise regularization, Pearson, *ρ*-sctransform and Spearman were calculated as 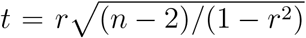 given the correlation estimate *r*, and two-sided *p*-values were evaluated under the standard normal distribution. The *p*-values for Normalisr were computed using the online code provided for its implementation.

### 4.3 Experiments with permuted scRNA-seq data

To generate null data sets from a given scRNA-seq data set with co-expression levels at or close to zero among all gene pairs while preserving gene expression levels, we first calculated normalized expression level for each gene *j* in cell *i*, written as *y_ij_* = *x_ij_*/*s_i_*. Then, for each gene *j*, we randomly permuted the normalized expressions 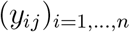 across *n* cells. After permutation, gene expressions were decorrelated and no gene pairs were expected to coexpress. Finally, the UMI count of gene *j* from cell *i* in the permuted data was calculated by sampling from 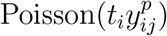, where 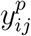 is the normalized expression level after permutation and *t_i_* is the desired sequencing depth in cell *i*. For the varying and constant sequencing depth settings in Figure 1, we set *t_i_* to the observed sequencing depth *s_i_* and median(*s*_1_, …, *s_n_*), respectively.

For numerical results in Figure 1 and Figure S2, we used the snRNA-seq data from Lau et al. (2020) and selected excitatory neurons from control subjects. The distribution of sequencing depths is long-tailed with a median of 5,833 (Figure S8). We randomly sampled 1,000 cells and sampled 500 genes from the top 5,000 highly expressed genes with probabilities proportional to the inverse density of expression levels. This ensures that the sampled genes could cover the range of expression levels. In the x-axis of Figure 1, we plotted expression levels at the scale of log_10_(*μ_j_*) + 3.

### 4.4 A simple illustration of the expression-level-dependent attenuation bias

To illustrate how errors from the Poisson measurement model in (1) can bias co-expression estimates, we conduct a short analysis under a much simplified case that directly calculates Pearson correlations of UMI counts. The analysis is similar to that in Wang et al. (2022), though *s_*i*_* was not considered there. From (1) and for genes *j*, *j*′, we have

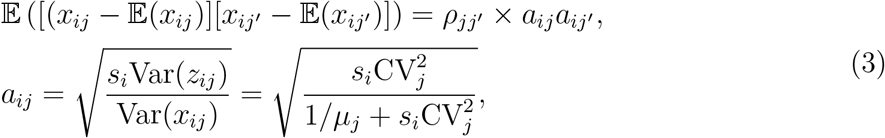

where CV*_j_* is the coefficient of variation of gene *j* defined as 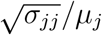. To measure the true correlation *ρ_jj′_*, the correlation based on UMI counts *x_ij_* and *x_ij′_* is always biased towards zero, as *a_ij_a_ij′_* < 1 when 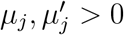. We refer to *a_ij_*, derived under the Poisson measurement model in (1), as the attenuation factor in this analysis.

When CV_*j*_’s are fixed, the attenuation factor *a_ij_* is closer to 1 for highly expressed genes with a larger *μ_j_*. Correspondingly, correlations are more accurately estimated for highly expressed genes and more attenuated for lowly expressed genes, assuming *s_i_*’s do not vary across cells. Based on a real snRNA-seq data set from Lau et al. (2020), we indeed observed that the estimated *a_ij_* approached 1 as the gene expression level increased (Figure S9). With *s_i_*’s varying across cells, the UMI counts for a pair of genes across cells are not identically distributed. In this case, it is difficult to analytically demonstrate the combined effect of the attenuation bias and the varying sequencing depths on co-expression estimation.

### 4.5 Simulating from the multivariate expression-measurement model

To simulate gene expression data from model (1), we combine a marginal negative binomial model and a copula-based approach that can simulate multivariate count data following a pre-specified co-expression matrix.

We specified the distribution of true expression level *z_ij_* to be Gamma(*α_j_*, *β_j_*) where *μ_j_* = *α_j_β_j_* and 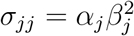 correspond to the marginal mean and variance in (1). Conditional on *z_ij_*, we simulated counts *x_ij_* from Poisson(*s_i_z_ij_*) independently for cell *i* and gene *j*. Marginally, this Poisson-Gamma mixture is equivalent to a negative binomial model on *x_ij_*, which is commonly used to model droplet-based single cell data (Risso et al., 2018; Hafemeister and Satija, 2019; Svensson, 2020; He et al., 2021). In our simulations, *μ_j_, σ_jj_* and *s_i_* are estimated or sampled from real data (see Section S2). Next, given a *p*×*p* correlation matrix *R*, we adopted a Gaussian copula to simulate correlated Gamma random variables (Tian et al., 2021; Sun et al., 2021). In particular, we first simulated samples (*v*_*i*1_, …, *v_ip_*) from a multivariate normal distribution with mean 0 and correlation *R* and then computed 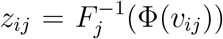, where Φ(·) is the cumulative distribution function (CDF) of a standard normal distribution and *F_j_*(·) is the CDF of Gamma(*α_j_, β_j_*). In Figure 2B, the matrix *R* was estimated from Lau et al. (2020) and in Figure 3A, the modular matrix *R* was generated from a network model. These details can be found in Section S2.

### 4.6 Differential co-expression analysis

For differential co-expression analysis, we first estimated co-expression networks from the disease and control groups separately. For the group with more cells, we randomly sampled a subset of cells such that the two groups had the same number of cells when estimating coexpressions. For each gene pair, we calculated the difference between co-expression estimates and assessed the statistical significance using a permutation test, where we randomly permuted the group labels 100 times and built a null distribution of differences in co-expressions. We then applied WGCNA (Langfelder and Horvath, 2008) to the significantly differentially co-expressed pairs (BH-adjusted *p*-values<0.05) with the soft-thresholding power set to 1 and extracted differentially co-expressed modules.

## 5 Data Availability

All data used in this work are publicly available, and the description, accession numbers and links of each data set are in Table S1. Codes that implement CS-CORE are covered by the MIT License and are available at GitHub: https://github.com/ChangSuBiostats/CS-CORE.

## 6 Acknowledgements

Zhang’s research is supported by NSF DMS-2015190 and DMS-2210469. Cai’s research is supported in part by DOD W81XWH2110019. Su, Xu, Shan, and Zhao’s research is supported in part by NIH R01 GM134005 and R56 AG074015.

