## supplementary_materials for "Cell-type-specific co-expression inference from single cell RNA-sequencing data"

#### S1 CS-CORE algorithm details

In CS-CORE, we estimate mean $\mu_j$, variance $\sigma_{jj}$ and covariance $\sigma_{jj'}$, respectively, via the following weighted least squares regressions:

$$\min_{\mu} \sum_i w_{ij}(x_{ij} - s_i\mu)^2,$$
$$\min_{\sigma} \sum_i h_{ij}[(x_{ij} - s_i\mu_j)^2 - s_i\mu_j - s_i^2\sigma]^2,$$
$$\min_{\sigma} \sum_i g_{ijj'}[(x_{ij} - s_i\mu_j)(x_{ij'} - s_i\mu_{j'}) - s_i^2\sigma]^2,$$

where $w_{ij}$, $h_{ij}$ and $g_{ijj'}$ are weights, respectively, defined as $w_{ij} = 1/(s_i\mu_j + s_i^2\sigma_{jj})$, $h_{ij} = w_{ij}^2$ and $g_{ijj'} = w_{ij}w_{ij'}$. Correspondingly, we have the following weighted least squares operators for estimating $\mu_j, \sigma_{jj}$, and $\sigma_{jj'}$, respectively,

$$f_{\mu_j}(w_{ij}) = \frac{\sum_i (s_i x_{ij}) w_{ij}}{\sum_i s_i^2 w_{ij}},$$
$$f_{\sigma_{jj}}(\mu_j, h_{ij}) = \frac{\sum_i s_i^2[(x_{ij} - s_i\mu_j)^2 - s_i\mu_j]h_{ij}}{\sum_i s_i^4 h_{ij}},$$
$$f_{\sigma_{jj'}}(\mu_j, \mu_{j'}, g_{ijj'}) = \frac{\sum_i s_i^2(x_{ij} - s_i\mu_j)(x_{ij'} - s_i\mu_{j'})g_{ijj'}}{\sum_i s_i^4 g_{ijj'}}.$$

In CS-CORE, we update the parameters $\mu_j, \sigma_{jj}$, and $\sigma_{jj'}$ and the weights used in their estimation via an IRLS procedure, where the weights are first initialized with OLS estimators of $\mu_j$'s and $\sigma_{jj}$'s. The detailed algorithm is presented in Algorithm S1.

#### S2 Details of simulating expression data from model (1)

In (1), we proposed the following expression-measurement model:

$$(z_{i1}, \ldots, z_{ip}) \sim F_p(\boldsymbol{\mu}, \boldsymbol{\Sigma}), \quad x_{ij}|z_{ij} \sim \text{Poisson}(s_i z_{ij}),$$

**Algorithm S1** CS-CORE estimation

**Input:** UMI count matrix $X = (x_{ij})_{n\times p}$ with $n$ cells and $p$ genes, sequencing depths $\{s_i\}_{i=1}^n$

Set $\Delta^{(0)} = 1$. Set $t = 0$.

// Estimate $\mu_j$'s and $\sigma_{jj}$'s

// Initialize with ordinary least sqaures

**for** $j = 1, \ldots, p$ **do**

  $\hat{\mu}_j^{(t)} \leftarrow f_{\mu_j}(1)$, $(\hat{\sigma}_{jj})^{(t)} \leftarrow f_{\sigma_{jj}}(\hat{\mu}_j^{(t)}, 1)$.

**end for**

// Iteratively reweighted least squares

**while** $\Delta^{(t)} \geq 0.05$ **do**

  $t \leftarrow t + 1$

  // Regularize $\theta_j$ estimates for weighting

  $\hat{\theta}^{(t)} \leftarrow \text{median}_j\{(\hat{\sigma}_{jj})^{(t-1)}/(\hat{\mu}_j^{(t-1)})^2\}$

  // Update $\mu_j, \sigma_{jj}$ estimates

  **for** $j = 1, \ldots, p$ **do**

    $w_{ij}^{(t)} \leftarrow 1/[s_i\hat{\mu}_j^{(t-1)} + s_i^2(\hat{\mu}_j^{(t-1)})^2 \times \hat{\theta}^{(t)}]$

    $\hat{\mu}_j^{(t)} \leftarrow f_{\mu_j}(w_{ij}^{(t)})$

    $h_{ij}^{(t)} \leftarrow 1/[s_i\hat{\mu}_j^{(t)} + s_i^2(\hat{\mu}_j^{(t)})^2 \times \hat{\theta}^{(t)}]^2$

    $(\hat{\sigma}_{jj})^{(t)} \leftarrow f_{\sigma_{jj}}(\hat{\mu}_j^{(t)}, h_{ij}^{(t)})$

  **end for**

  // Assess convergence

  $\Delta^{(t)} \leftarrow \max_j |\log(\hat{\sigma}_{jj})^{(t)} - \log(\hat{\sigma}_{jj})^{(t-1)}|$

**end while**

// Estimate $\sigma_{jj'}$'s

Let $\hat{\theta} = \hat{\theta}^{(t)}$, $\hat{\mu}_j = \hat{\mu}_j^{(t)}, \hat{\sigma}_{jj} = (\hat{\sigma}_{jj})^{(t)}$ for $j, j' = 1, \ldots, p$.

**for** $j, j' = 1, \ldots, p, j \neq j'$ **do**

  $g_{ijj'} = 1/\{[s_i\hat{\mu}_j + s_i^2(\hat{\mu}_j)^2 \times \hat{\theta}][s_i\hat{\mu}_{j'} + s_i^2(\hat{\mu}_{j'})^2 \times \hat{\theta}]\}$

  $\hat{\sigma}_{jj'} = f_{\sigma_{jj'}}(\hat{\mu}_j, \hat{\mu}_{j'}, g_{ijj'})$

  $\hat{\rho}_{jj'} = \hat{\sigma}_{jj'}/\sqrt{\hat{\sigma}_{jj}\hat{\sigma}_{j'j'}}$

**end for**

**Output:** $\hat{\mu}_j, \hat{\sigma}_{jj}, \hat{\sigma}_{jj'}$ for $j, j' = 1, \ldots, p$

where $F_p(\boldsymbol{\mu}, \boldsymbol{\Sigma})$ is a nonnegative $p$-variate distribution with mean vector $\boldsymbol{\mu} = (\mu_1, \ldots, \mu_p)$ and covariance matrix $\boldsymbol{\Sigma} = (\sigma_{jj'})_{p \times p}$. For experiments in Figure 2B-C and 3A, we used the copula method described in Section 4.5 to simulate from the above model. Next, we discuss how the parameters were set in each simulation setting.

We generated $n = 5,000$ cells on $p = 500$ genes in Figure 2B-C. To select values of $\mu_j, \sigma_{jj}$ and $s_i$ that resemble real data, we utilized the single cell data on excitatory neurons from Lau et al. (2020). Specifically, for sequencing depths $s_i$, we sampled from the observed values in real data. For mean and variance parameters $\mu_j$ and $\sigma_{jj}$, we fitted a negative binomial generalized linear model (Ahlmann-Eltze and Huber, 2020) that estimated $\mu_j$ and $\text{CV}_j^2 = \sigma_{jj}/\mu_j^2$ across genes using maximum likelihood. We then set $\mu_j, j = 1, \ldots, 500$ in simulation to be the estimates for the top 500 highly expressed genes. As estimates of $\text{CV}_j$'s from maximum likelihood estimation can be highly variable (Lause et al., 2021), we exploited a mean-dispersion trend that is well-known in bulk RNA-seq data (Robinson et al., 2010; McCarthy et al., 2012; Wu et al., 2013; Love et al., 2014) and recently also observed in single-cell data (Hafemeister and Satija, 2019; Choudhary and Satija, 2022) by fitting $\log_{10}(\text{CV}_j^2)$ as a function of $\log_{10}(\mu_j)$ with kernel smoothing regression (Hafemeister and Satija, 2019), denoted as $\hat{f}_j(\cdot)$. Finally, we used estimates of $\mu_j$ and $\sigma_{jj} = 10^{\hat{f}_j(\mu_j)} \times \mu_j^2$ as the parameters in the marginal Gamma distributions. To specify a matrix $R$ that resembles real data in Figure 2B, we first applied CS-CORE to estimate the co-expression matrix for the top 500 highly expressed genes in excitatory neurons from Lau et al. (2020) and only focused on significantly co-expressed gene pairs (BH-adjusted $p$-values $< 0.05$). We further set co-expression estimates with absolute values less than 0.5 to 0 to encourage sparsity. Finally, we scaled off-diagonal entries by 3.1 such that the scaled matrix was positive definite.

Next, we generated $n = 2,000$ cells on $p = 100$ genes in Figure 3A. To select values of $\mu_j, \sigma_{jj}$ and $s_i$ that resemble real data, we also utilized the single cell data on excitatory neurons from Lau et al. (2020). For sequencing depths $s_i$, we sampled from the observed values in real data. We then estimated $\mu_j$ and $\sigma_{jj}$ for 100 genes that were randomly selected from the top 2,000 highly expressed genes using a procedure similar as above. To generate a co-expression matrix $R$ with the clustering structure in Figure 3A, we adopted the degree-corrected Stochastic Block model (Lee and Wilkinson, 2019) and used `BlockModel.Gen` in R package *randnet* (v.0.1) to simulate a binary network of size 100, where we specified four co-expressed modules and set the ratio of within-block edges and between-block edges to be 1,000. We further set the node degree (i.e. sum of co-expressed genes) distribution to be a power-law distribution with power equal to 4 and mean equal to 20. Given the binary network, we then simulated co-expression values from a truncated standard normal distribution between 0.9 and 1. Finally, we used the software from Sun and Vandenberghe

(2015) to obtain a closest positive definite correlation matrix and removed five genes that had minimal connections with other genes in their assigned clusters.

The software for Normalisr, Noise regularization, propr and sctransform require the entire gene count matrix as input, for example, to calculate sequencing depth per cell. In this case, we further simulated independent counts with marginal statistics resembling real data for genes that were not included in Figure 2B-C and 3A.

### S3 Details of enrichment and reproducibility analyses

#### S3.1 GO and Reactome Enrichment analysis

Given a gene set identified via clustering analysis, for enrichment in biological pathways, we performed enrichment analysis of Gene Ontology (GO) using R package clusterProfiler (v.4.2.2) from Wu et al. (2021) and Reactome Pathway Database (Reactome) using R package ReactomePA (v.1.38.0) from Yu and He (2016) with background genes set to all genes that were used in the clustering analysis. We reported significantly enriched GO terms which had BH-adjusted $p$-values less than 0.05.

#### S3.2 Overlap with known TF-target gene pairs

Given a given scRNA-seq data set and a co-expression estimation method, for assessing whether the method can identify known TF-target gene pairs, we took the union of the top 5,000 highly expressed genes and genes in the TRRUST database (Han et al., 2018), calculated the $p$-values in testing for co-expressed gene pairs using either CS-CORE or $\rho$-sctransform and counted the number of significant gene pairs that overlapped with the TRRUST database across different $p$-value cutoffs.

#### S3.3 Reproducibility analysis

To evaluate the reproducibility of gene pairs identified with CS-CORE and $\rho$-sctransform in five brain cell types (Figure 4A and Figure S6, S7), we counted the number of reproduced pairs across different $p$-value cutoffs between two sets of independent snRNA-seq data sets on the brain. Specifically, for each cell type, we used the cells on control subjects from Lau et al. (2020) and Mathys et al. (2019), took the intersection of top 5,000 highly expressed genes between these two data sets, computed co-expression $p$-values for the intersected genes and compared significant gene pairs inferred in these two data sets (Figure 4A). We also used the cells on control subjects from Lau et al. (2020) and Morabito et al. (2021) and compared significant gene pairs similarly (Figure S6). To evaluate the reproducibility in five immune cell types in PBMC (Figure S7), we used cells of control subjects from two independent PBMC data sets Wilk et al. (2020) and Unterman et al. (2022) and counted the number of reproduced gene pairs in each cell type following a similar procedure.

#### S4 Sequencing depth variations cannot be adjusted by standard marginal normalization

We conduct a simple experiment and demonstrate that standard marginal normalization, such as scaling or log-normalization, cannot remove the confounding effect of varying sequencing depths on inferring co-expressions. In this experiment, we simulated independent UMI counts as in Figure S2 and focused on a gene pair that had expression levels ranked 362 and 500 among 28,412 genes. We computed the scaled data as $x_i/s_i$ and the log normalized data as $\log(x_i/s_i \times 10^4 + 1)$, and show the scatter plots based on original, scaled and log normalized counts in Figure S1.

Given two integers $a$ and $b$, all cells with UMI counts $a, b$ for these two genes, respectively, are plotted to the point $(a, b)$ in the original UMI counts scatter plot. Interestingly, cells at this point, turning into $(a/s_i, b/s_i)$ after scaling, will be stretched out to form a line with a slope $b/a$ and an intercept 0 in the scaled data, and turning into $(\log(a)-\log(s_i/10^4), \log(b)-\log(s_i/10^4))$ after log normalization, approximately forming a line with a slope 1 and an intercept $\log(b) - \log(a)$ in the log normalized data (Figure S1). These lines are artifacts of the normalization and can seriously inflate false positives when inferring co-expressions. For example, while expression data for these two genes were simulated independently, we got the following correlation estimates along with $p$-values (only calculated for methods that offer tests) for existing methods that use log-normalized data: locCSN=0.61, Pearson=0.14 ($p < 0.05$), propr=0.50, Spearman=0.07 ($p < 0.05$) and SpQN=0.14.

#### S5 Data summary and pre-processing

A summary of the data sets analyzed in our work is given in Table S1. For cell type labels of the cells, we used the cell type labels provided by authors of Mathys et al. (2019), Morabito et al. (2021), Wilk et al. (2020), and Unterman et al. (2022). Cell type labels were not provided for the data set from Lau et al. (2020) and we annotated the cell types following the procedure described in Lau et al. (2020).

To conduct the reproduciblity analysis in five major immune cell types between Wilk

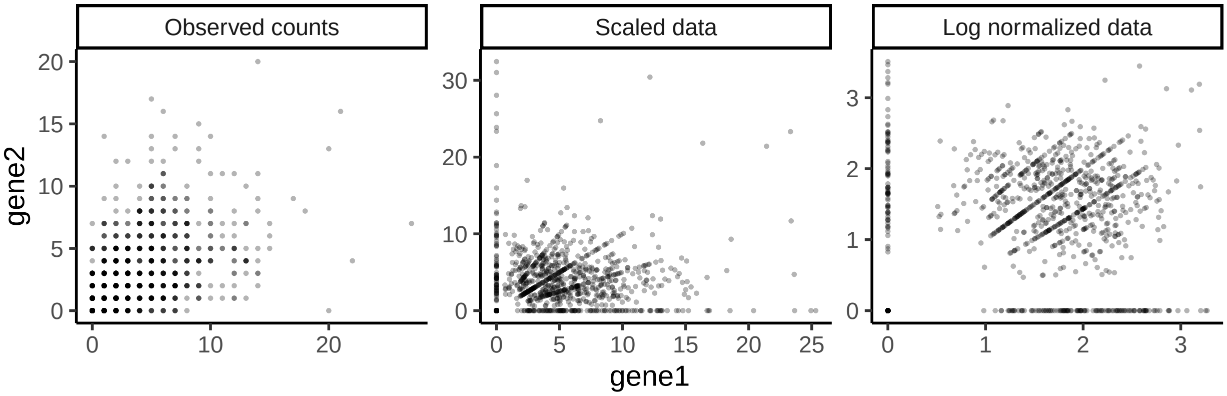


Figure S1: Expressions of an independent gene pair in raw UMI counts, scaled and log-normalized counts. We computed the scaled data as $x_i/s_i$ and the log normalized data as $\log(x_i/s_i \times 10^4 + 1)$.

et al. (2020) and Unterman et al. (2022), we combined the Naive B cells and Memory B cells from Unterman et al. (2022) to compare with B cells from Wilk et al. (2020); we combined the CD14 Monocytes and CD16 Monocytes from Wilk et al. (2020) to compare with the combination of classical monocytes and NC and IM monocytes from Unterman et al. (2022); we combined the Naive CD4 T, Memory CD4 T and Memory CD4 and MAI T cells from Unterman et al. (2022) to compare with CD4 T cells from Wilk et al. (2020); we combined the Memory CD8 T, Naive CD8 T, Effector T and IFN-activated CD8 T cells from Unterman et al. (2022) to compare with CD8 T cells from Wilk et al. (2020); we combined the NK CD56dim and NK CD56bright cells from Unterman et al. (2022) to compare with the NK cells from Wilk et al. (2020).

Table S1: Summary of single-cell (nucleus) RNA-seq data used in analyses

| Data sets | Lau et al. (2020) | Mathys et al. (2019) | Morabito et al. (2021) | Wilk et al. (2020) | Unterman et al. (2022) |
|---|---|---|---|---|---|
| Tissue | Brain | Brain | Brain | PBMC | PBMC |
| Data | snRNA-seq | snRNA-seq | snRNA-seq | scRNA-seq | scRNA-seq |
| #cells/nucleus | 169,500 | 70,634 | 61,472 | 44,721 | 153,554 |
| #cell types | 6 | 8 | 7 | 13 | 29 |
| Median seq depth | 2,600 | 1,474 | 6,382 | 1,946 | 3,618 |
| #genes | 28,412 | 17,926 | 36,114 | 26,361 | 33,538 |
| #samples | 21 | 48 | 18 | 14 | 31 |
| Data availability | GSE157827 | syn21261143 | syn22079621 | Website, 'Peripheral Blood Mononuclear Cells (PBMCs)' | GSE155224 |

### S6 GO enrichment analysis of modules identified with CS-CORE and $\rho$-sctransform

In Tables S2-S3, we estimated co-expressions using CS-CORE and $\rho$-sctransform, respectively, using cells from control subjects in Lau et al. (2020) for five major brain cell types, including astrocyte, excitatory neuron, inhibitory neuron, oligodendrocyte, and microglia, and performed enrichment analyses of extracted co-expression modules. Specifically, for each cell type, we performed clustering analyses following WGCNA (Langfelder and Horvath, 2008) with the soft-thresholding power set to 1 to obtain co-expressed gene modules. We conducted GO enrichment analyses on the identified modules (see Section S3.1) and the top three GO terms with the strongest enrichment signals are presented in Tables S2-S3 for modules with at least one highly significant GO term (BH-adjusted $p$-values $< 0.001$).

Table S2: Gene Ontology enrichment analysis of co-expressed gene modules identified with CS-CORE using brain samples from Lau et al. (2020). Representative GO terms capturing cell-type-specific biological functions are highlighted in blue.

| Cell type | module | GO term ID | GO term | Adjusted $p$-value | Gene ratio |
|---|---|---|---|---|---|
| Astrocyte | 1 | **GO:0099536** | **synaptic signaling** | **5.14e-10** | **56/333** |
| | | GO:0007268 | chemical synaptic transmission | 5.14e-10 | 54/333 |
| | | GO:0098916 | anterograde trans-synaptic signaling | 5.14e-10 | 54/333 |
| | 2 | **GO:0006457** | **protein folding** | **1.26e-08** | **17/124** |
| | | GO:0061077 | chaperone-mediated protein folding | 1.52e-05 | 9/124 |
| | | GO:0034605 | cellular response to heat | 3.19e-05 | 9/124 |
| | 3 | GO:0002181 | cytoplasmic translation | 1.27e-14 | 18/81 |
| | | GO:0006119 | oxidative phosphorylation | 3.37e-09 | 12/81 |
| | | GO:0009060 | aerobic respiration | 2.33e-07 | 12/81 |
| | 4 | **GO:0071456** | **cellular response to hypoxia** | **5.26e-09** | **8/16** |
| | | GO:0036294 | cellular response to decreased oxygen levels | 5.34e-09 | 8/16 |
| | | GO:0071453 | cellular response to oxygen levels | 7.70e-09 | 8/16 |
| | 5 | GO:0016579 | protein deubiquitination | 6.33e-04 | 4/10 |
| | | GO:0070646 | protein modification by small protein removal | 6.33e-04 | 4/10 |
| | | GO:0071108 | protein K48-linked deubiquitination | 4.79e-02 | 2/10 |
| Inhibitory neuron | 1 | GO:0002181 | cytoplasmic translation | 2.07e-12 | 51/1468 |
| | | GO:0009060 | aerobic respiration | 2.30e-09 | 44/1468 |
| | | GO:0006119 | oxidative phosphorylation | 5.93e-08 | 33/1468 |
| | 2 | **GO:0097060** | **synaptic membrane** | **6.39e-06** | **111/1469** |
| | | GO:0045211 | postsynaptic membrane | 9.10e-05 | 78/1469 |
| | | GO:0098590 | plasma membrane region | 2.60e-03 | 190/1469 |
| Oligodendrocyte | 1 | GO:0002181 | cytoplasmic translation | 4.27e-14 | 66/1343 |
| | | GO:0006412 | translation | 4.82e-07 | 126/1343 |
| | | GO:0043043 | peptide biosynthetic process | 5.88e-07 | 128/1343 |
| | 2 | **GO:0099536** | **synaptic signaling** | **4.15e-12** | **77/557** |
| | | GO:0007268 | chemical synaptic transmission | 4.15e-12 | 74/557 |
| | | GO:0098916 | anterograde trans-synaptic signaling | 4.15e-12 | 74/557 |
| | 3 | **GO:0008203** | **cholesterol metabolic process** | **2.00e-08** | **6/10** |
| | | GO:1902652 | secondary alcohol metabolic process | 2.00e-08 | 6/10 |
| | | GO:0016125 | sterol metabolic process | 2.00e-08 | 6/10 |
| Microglia | 1 | **GO:0006952** | **defense response** | **3.54e-08** | **300/2340** |
| | | GO:0098542 | defense response to other organism | 2.61e-07 | 211/2340 |
| | | GO:0007155 | cell adhesion | 4.56e-07 | 302/2340 |
| | 2 | **GO:0007268** | **chemical synaptic transmission** | **1.72e-06** | **53/428** |
| | | GO:0098916 | anterograde trans-synaptic signaling | 1.72e-06 | 53/428 |
| | | GO:0099537 | trans-synaptic signaling | 2.20e-06 | 53/428 |
| | 3 | **GO:0002181** | **cytoplasmic translation** | **1.95e-69** | **68/193** |
| | | GO:0006412 | translation | 2.70e-51 | 84/193 |
| | | GO:0043043 | peptide biosynthetic process | 9.70e-51 | 84/193 |
| | 4 | **GO:0006457** | **protein folding** | **5.16e-25** | **25/69** |
| | | GO:0061077 | chaperone-mediated protein folding | 2.42e-14 | 13/69 |
| | | GO:0006458 | ‘de novo’ protein folding | 9.10e-12 | 10/69 |

Table S3: Gene Ontology enrichment analysis of co-expressed gene modules identified with $\rho$-sctransform using brain samples from Lau et al. (2020). Representative GO terms that capture cell-type-specific biological functions are highlighted in blue.

| Cell type | module | GO term ID | GO term | Adjusted $p$-value | Gene ratio |
|---|---|---|---|---|---|
| Astrocyte | 1 | GO:0030141 | secretory granule | 3.96e-05 | 61/629 |
| | | GO:0005773 | vacuole | 9.01e-05 | 66/629 |
| | | GO:0000323 | lytic vacuole | 9.01e-05 | 60/629 |
| | 2 | **GO:0099536** | **synaptic signaling** | **2.02e-07** | **47/287** |
| | | GO:0007268 | chemical synaptic transmission | 2.02e-07 | 45/287 |
| | | GO:0098916 | anterograde trans-synaptic signaling | 2.02e-07 | 45/287 |
| | 3 | **GO:0034765** | **regulation of ion transmembrane transport** | **1.08e-04** | **10/39** |
| | | GO:0043269 | regulation of ion transport | 1.46e-04 | 11/39 |
| | | GO:0034762 | regulation of transmembrane transport | 3.06e-04 | 10/39 |
| | 4 | GO:0002181 | cytoplasmic translation | 3.15e-09 | 10/31 |
| | | GO:0006412 | translation | 1.93e-05 | 11/31 |
| | | GO:0009060 | aerobic respiration | 1.93e-05 | 7/31 |
| Excitatory neuron | 1 | GO:0002181 | cytoplasmic translation | 2.42e-07 | 42/1648 |
| | | GO:0043043 | peptide biosynthetic process | 1.12e-05 | 115/1648 |
| | | GO:0006412 | translation | 1.30e-05 | 112/1648 |
| Inhibitory neuron | 1 | GO:0002181 | cytoplasmic translation | 8.88e-14 | 38/661 |
| | | GO:0006518 | peptide metabolic process | 3.08e-11 | 89/661 |
| | | GO:0043603 | cellular amide metabolic process | 6.87e-11 | 101/661 |
| | 2 | GO:0002181 | cytoplasmic translation | 9.34e-05 | 7/30 |
| | | GO:0006412 | translation | 1.80e-02 | 8/30 |
| | | GO:0043043 | peptide biosynthetic process | 1.80e-02 | 8/30 |
| Oligodendrocyte | 1 | GO:0002181 | cytoplasmic translation | 1.07e-17 | 66/1167 |
| | | GO:0006412 | translation | 1.44e-10 | 123/1167 |
| | | GO:0043043 | peptide biosynthetic process | 1.10e-09 | 123/1167 |
| | 2 | **GO:0007268** | **chemical synaptic transmission** | **4.44e-11** | **69/518** |
| | | GO:0098916 | anterograde trans-synaptic signaling | 4.44e-11 | 69/518 |
| | | GO:0099536 | synaptic signaling | 4.44e-11 | 71/518 |
| | 3 | GO:0045211 | postsynaptic membrane | 3.36e-05 | 9/45 |
| | | GO:0031226 | intrinsic component of plasma membrane | 2.42e-04 | 13/45 |
| | | GO:0005887 | integral component of plasma membrane | 3.89e-04 | 12/45 |
| Microglia | 1 | GO:0007155 | cell adhesion | 4.82e-13 | 365/2807 |
| | | GO:0022610 | biological adhesion | 4.82e-13 | 366/2807 |
| | | GO:0098609 | cell-cell adhesion | 1.08e-07 | 223/2807 |
| | 2 | **GO:0006457** | **protein folding** | **4.13e-14** | **33/344** |
| | | GO:0061077 | chaperone-mediated protein folding | 2.32e-08 | 16/344 |
| | | GO:0042026 | protein refolding | 3.19e-07 | 10/344 |
| | 3 | GO:0002181 | cytoplasmic translation | 5.11e-61 | 70/271 |
| | | GO:0006412 | translation | 2.09e-44 | 91/271 |
| | | GO:0043043 | peptide biosynthetic process | 8.28e-44 | 91/271 |

In Tables S4-S5, we evaluated the differential co-expression network estimated using CS-CORE and $\rho$-sctranform (see Section 4.6), respectively, using microglia cells from control subjects and AD patients. We performed clustering analyses following WGCNA (Langfelder and Horvath, 2008) with the soft-thresholding power set to 1 to obtain differentially co-expressed gene modules. We then conducted GO enrichment analyses (see Section S3.1) on the identified modules and the top three GO terms with the strongest enrichment signals are presented in Tables S4-S5 for modules with at least one significant GO term (BH-adjusted $p$-values $< 0.05$). For results from $\rho$-sctranform in Table S5, we did not identify any GO term that captures cell-type-specific biological functions or cell-type-specific disease-related biological pathways in the presented list.

Table S4: Gene Ontology enrichment analysis of differentially co-expressed gene modules identified with CS-CORE in microglia using brain samples from Lau et al. (2020). Representative GO terms capturing cell-type-specific biological functions and/or cell-type-specific disease-related biological pathways are highlighted in blue.

| Cell type | Module | GO term ID | GO term | Adjusted $p$-value | Gene ratio |
|---|---|---|---|---|---|
| Microglia | 1 | **GO:0006457** | **protein folding** | **1.27e-03** | **8/65** |
| | | GO:0006986 | response to unfolded protein | 5.64e-03 | 7/65 |
| | | GO:0035966 | response to topologically incorrect protein | 1.26e-02 | 7/65 |
| | 2 | GO:0031224 | intrinsic component of membrane | 1.84e-04 | 32/67 |
| | | **GO:0098978** | **glutamatergic synapse** | **1.03e-03** | **11/67** |
| | | GO:0016021 | integral component of membrane | 1.03e-03 | 29/67 |
| | 3 | GO:0002181 | cytoplasmic translation | 4.63e-16 | 14/30 |
| | | GO:0006412 | translation | 3.31e-10 | 15/30 |
| | | GO:0043043 | peptide biosynthetic process | 3.31e-10 | 15/30 |
| | 4 | GO:0051171 | regulation of nitrogen compound metabolic process | 4.59e-02 | 11/11 |
| | | GO:0080090 | regulation of primary metabolic process | 4.59e-02 | 11/11 |
| | | **GO:0019901** | **protein kinase binding** | **4.19e-03** | **6/11** |

Table S5: Gene Ontology enrichment analysis of differentially co-expressed gene modules identified with $\rho$-sctransform in microglia using brain samples from Lau et al. (2020). We did not identify any GO term that captures cell-type-specific biological functions or cell-type-specific disease-related biological pathways in the presented list.

| Cell type | Module | GO term ID | GO term | Adjusted $p$-value | Gene ratio |
|---|---|---|---|---|---|
| Microglia | 1 | GO:0005887 | integral component of plasma membrane | 4.03e-09 | 32/114 |
| | | GO:0031226 | intrinsic component of plasma membrane | 9.98e-09 | 32/114 |
| | | GO:0031224 | intrinsic component of membrane | 2.99e-04 | 45/114 |
| | 2 | GO:0035148 | tube formation | 3.25e-02 | 6/68 |
| | 3 | GO:0002181 | cytoplasmic translation | 1.85e-18 | 16/38 |
| | | GO:0006412 | translation | 7.79e-11 | 17/38 |
| | | GO:0043043 | peptide biosynthetic process | 7.79e-11 | 17/38 |
| | 4 | GO:0009897 | external side of plasma membrane | 3.01e-02 | 4/14 |
| | | GO:0043235 | receptor complex | 3.35e-02 | 4/14 |
| | | GO:0099503 | secretory vesicle | 3.35e-02 | 6/14 |

In Tables S6-S8, we evaluated CS-CORE using cells from control subjects in Wilk et al. (2020) for five major immune cell types, including B cells, CD4 positive T cells, CD8 positive T cells, monocytes and natural killer (NK) cells, and performed enrichment analyses of extracted co-expression modules. Specifically, for each cell type, we performed clustering analyses following WGCNA (Langfelder and Horvath, 2008) with the soft-thresholding power set to 1 to obtain co-expressed gene modules. We then conducted GO enrichment analyses on the identified modules and the top three GO terms with the strongest enrichment signals are presented in Tables S6-S8 for modules with at least one highly significant GO term (BH-adjusted $p$-values $< 0.001$). The same procedure was applied to $\rho$-sctransform, and the results are given in Tables S9-S10.

Table S6: Gene Ontology enrichment analysis of cell-type-specific co-expressed gene modules identified with CS-CORE in B cells and CD4 T cells using PBMC samples from Wilk et al. (2020). Representative GO terms that capture cell-type-specific biological functions are highlighted in blue.

| Cell type | Module | GO term ID | GO term | Adjusted *p*-value | Gene ratio |
|---|---|---|---|---|---|
| B | 1 | GO:0002181 | cytoplasmic translation | 1.29e-80 | 83/232 |
| | | GO:0006412 | translation | 3.12e-43 | 94/232 |
| | | GO:0043043 | peptide biosynthetic process | 3.12e-43 | 94/232 |
| | 2 | GO:0030029 | actin filament-based process | 2.85e-04 | 21/107 |
| | | GO:0030036 | actin cytoskeleton organization | 2.85e-04 | 20/107 |
| | | GO:0007015 | actin filament organization | 4.36e-04 | 16/107 |
| | 3 | **GO:0019886** | **antigen processing and presentation of exogenous peptide antigen via MHC class II** | **7.77e-16** | **14/109** |
| | | GO:0002495 | antigen processing and presentation of peptide antigen via MHC class II | 1.04e-15 | 14/109 |
| | | GO:0002399 | MHC class II protein complex assembly | 1.04e-15 | 12/109 |
| | 4 | **GO:0002250** | **adaptive immune response** | **4.97e-05** | **13/37** |
| | | GO:0006910 | phagocytosis, recognition | 4.11e-04 | 6/37 |
| | | GO:0042113 | B cell activation | 4.11e-04 | 10/37 |
| | 5 | **GO:0035455** | **response to interferon-alpha** | **1.66e-04** | **4/25** |
| | | GO:0009615 | response to virus | 4.72e-04 | 8/25 |
| | | GO:0051607 | defense response to virus | 4.72e-04 | 7/25 |
| CD4T | 1 | GO:0002181 | cytoplasmic translation | 2.86e-60 | 86/425 |
| | | GO:0006412 | translation | 2.22e-28 | 108/425 |
| | | GO:0043043 | peptide biosynthetic process | 2.61e-28 | 108/425 |
| | 2 | GO:0070161 | anchoring junction | 1.77e-04 | 47/348 |
| | | GO:0015629 | actin cytoskeleton | 1.77e-04 | 29/348 |
| | | GO:0005911 | cell-cell junction | 6.65e-04 | 25/348 |
| | 3 | **GO:0051607** | **defense response to virus** | **5.96e-16** | **20/55** |
| | | GO:0140546 | defense response to symbiont | 5.96e-16 | 20/55 |
| | | GO:0009615 | response to virus | 8.52e-16 | 22/55 |
| | 4 | **GO:0002478** | **antigen processing and presentation of exogenous peptide antigen** | **7.25e-04** | **4/21** |
| | | GO:0019884 | antigen processing and presentation of exogenous antigen | 9.00e-04 | 4/21 |
| | | GO:0048002 | antigen processing and presentation of peptide antigen | 1.73e-03 | 4/21 |

Table S7: Gene Ontology enrichment analysis of cell-type-specific co-expressed gene modules identified with CS-CORE in CD8 T cells using PBMC samples from Wilk et al. (2020). Representative GO terms that capture cell-type-specific biological functions are highlighted in blue.

| Cell type | Module | GO term ID | GO term | Adjusted $p$-value | Gene ratio |
| --- | --- | --- | --- | --- | --- |
| CD8T | 1 | GO:0009060 | aerobic respiration | 3.82e-05 | 30/519 |
| | | GO:0006091 | generation of precursor metabolites and energy | 3.82e-05 | 46/519 |
| | | GO:0046034 | ATP metabolic process | 3.82e-05 | 34/519 |
| | 2 | GO:0006260 | DNA replication | 8.15e-15 | 50/545 |
| | | GO:0006259 | DNA metabolic process | 1.43e-10 | 93/545 |
| | | GO:0006281 | DNA repair | 2.05e-10 | 65/545 |
| | 3 | GO:0051301 | cell division | 1.56e-06 | 56/456 |
| | | GO:0033045 | regulation of sister chromatid segregation | 5.23e-05 | 16/456 |
| | | GO:0007088 | regulation of mitotic nuclear division | 5.23e-05 | 17/456 |
| | 4 | GO:0002181 | cytoplasmic translation | 6.91e-69 | 85/326 |
| | | GO:0043043 | peptide biosynthetic process | 2.63e-33 | 100/326 |
| | | GO:0006412 | translation | 3.91e-33 | 99/326 |
| | 5 | GO:0030036 | actin cytoskeleton organization | 6.45e-06 | 34/228 |
| | | GO:0032970 | regulation of actin filament-based process | 6.45e-06 | 25/228 |
| | | GO:0030029 | actin filament-based process | 6.45e-06 | 35/228 |
| | 6 | **GO:0042269** | **regulation of natural killer cell mediated cytotoxicity** | **7.24e-06** | **10/197** |
| | | GO:0002715 | regulation of natural killer cell mediated immunity | 7.24e-06 | 10/197 |
| | | GO:0001910 | regulation of leukocyte mediated cytotoxicity | 8.70e-06 | 12/197 |
| | 7 | GO:0040011 | locomotion | 5.24e-05 | 43/189 |
| | | GO:0048870 | cell motility | 5.15e-04 | 38/189 |
| | | GO:0051674 | localization of cell | 5.15e-04 | 38/189 |
| | 8 | GO:0062023 | collagen-containing extracellular matrix | 5.40e-06 | 7/46 |
| | | GO:0030312 | external encapsulating structure | 5.40e-06 | 7/46 |
| | | GO:0031012 | extracellular matrix | 5.40e-06 | 7/46 |
| | 9 | GO:0009612 | response to mechanical stimulus | 2.04e-05 | 8/39 |
| | | GO:0009628 | response to abiotic stimulus | 7.80e-05 | 15/39 |
| | | GO:0014070 | response to organic cyclic compound | 1.76e-03 | 12/39 |
| | 10 | GO:0001725 | stress fiber | 1.37e-06 | 5/24 |
| | | GO:0097517 | contractile actin filament bundle | 1.37e-06 | 5/24 |
| | | GO:0032432 | actin filament bundle | 1.37e-06 | 5/24 |
| | 11 | **GO:0034976** | **response to endoplasmic reticulum stress** | **1.81e-06** | **9/20** |
| | | GO:0006457 | protein folding | 5.52e-05 | 7/20 |
| | | GO:0036503 | ERAD pathway | 1.17e-03 | 5/20 |
| | 12 | GO:0016973 | poly(A)+ mRNA export from nucleus | 4.10e-04 | 3/11 |
| | | GO:0006406 | mRNA export from nucleus | 9.80e-03 | 3/11 |
| | | GO:0071427 | mRNA-containing ribonucleoprotein complex export from nucleus | 9.80e-03 | 3/11 |

Table S8: Gene Ontology enrichment analysis of cell-type-specific co-expressed gene modules identified with CS-CORE in monocytes (Mono) and natural killer (NK) cells using PBMC samples from Wilk et al. (2020). Representative GO terms that capture cell-type-specific biological functions are highlighted in blue.

| Cell type | Module | GO term ID | GO term | Adjusted *p*-value | Gene ratio |
|---|---|---|---|---|---|
| Mono | 1 | GO:0030036 | actin cytoskeleton organization | 9.03e-08 | 89/829 |
| | | GO:0030029 | actin filament-based process | 9.03e-08 | 94/829 |
| | | GO:0097435 | supramolecular fiber organization | 2.88e-07 | 90/829 |
| | 2 | GO:0002181 | cytoplasmic translation | 7.25e-65 | 89/437 |
| | | GO:0043043 | peptide biosynthetic process | 1.30e-34 | 114/437 |
| | | GO:0006412 | translation | 1.38e-34 | 113/437 |
| | 3 | **GO:0002399** | **MHC class II protein complex assembly** | **7.18e-14** | **10/90** |
| | | GO:0002503 | peptide antigen assembly with MHC class II protein complex | 7.18e-14 | 10/90 |
| | | GO:0002501 | peptide antigen assembly with MHC protein complex | 1.20e-12 | 10/90 |
| | 4 | **GO:0051607** | **defense response to virus** | **9.92e-28** | **33/85** |
| | | GO:0140546 | defense response to symbiont | 9.92e-28 | 33/85 |
| | | GO:0009615 | response to virus | 3.16e-24 | 34/85 |
| | 5 | **GO:1900117** | **regulation of execution phase of apoptosis** | **6.75e-06** | **5/36** |
| | | GO:2000272 | negative regulation of signaling receptor activity | 2.62e-05 | 5/36 |
| | | **GO:0019885** | **antigen processing and presentation of endogenous peptide antigen via MHC class I** | **8.50e-04** | **4/36** |
| | 6 | **GO:0006910** | **phagocytosis, recognition** | **9.30e-09** | **5/11** |
| | | GO:0006958 | complement activation, classical pathway | 1.08e-08 | 5/11 |
| | | GO:0002455 | humoral immune response mediated by circulating immunoglobulin | 2.78e-08 | 5/11 |
| NK | 1 | GO:0002181 | cytoplasmic translation | 9.20e-44 | 89/710 |
| | | GO:0006412 | translation | 1.01e-15 | 116/710 |
| | | GO:0043043 | peptide biosynthetic process | 2.02e-15 | 116/710 |
| | 2 | **GO:0019885** | **antigen processing and presentation of endogenous peptide antigen via MHC class I** | **2.47e-06** | **9/246** |
| | | GO:0002483 | antigen processing and presentation of endogenous peptide antigen | 3.21e-06 | 9/246 |
| | | GO:0001914 | regulation of T cell mediated cytotoxicity | 3.21e-06 | 10/246 |
| | 3 | **GO:0042110** | **T cell activation** | **9.82e-04** | **20/132** |
| | | GO:0001775 | cell activation | 9.82e-04 | 29/132 |
| | | GO:0045321 | leukocyte activation | 9.82e-04 | 27/132 |
| | 4 | GO:0030029 | actin filament-based process | 3.09e-15 | 34/104 |
| | | GO:0030036 | actin cytoskeleton organization | 3.09e-15 | 33/104 |
| | | GO:0007015 | actin filament organization | 1.17e-13 | 26/104 |
| | 5 | **GO:0051607** | **defense response to virus** | **1.08e-06** | **13/57** |
| | | GO:0140546 | defense response to symbiont | 1.08e-06 | 13/57 |
| | | GO:0045071 | negative regulation of viral genome replication | 1.08e-06 | 8/57 |
| | 6 | GO:0006457 | protein folding | 1.87e-08 | 14/60 |
| | | **GO:0034976** | **response to endoplasmic reticulum stress** | **1.06e-06** | **14/60** |
| | | GO:0061077 | chaperone-mediated protein folding | 4.16e-02 | 5/60 |
| | 7 | GO:0022616 | DNA strand elongation | 9.25e-04 | 4/43 |
| | | GO:0006270 | DNA replication initiation | 9.25e-04 | 4/43 |
| | | GO:0033260 | nuclear DNA replication | 9.25e-04 | 4/43 |
| | 8 | **GO:0019886** | **antigen processing and presentation of exogenous peptide antigen via MHC class II** | **8.60e-04** | **3/13** |
| | | GO:0002495 | antigen processing and presentation of peptide antigen via MHC class II | 8.60e-04 | 3/13 |
| | | GO:0002504 | antigen processing and presentation of peptide or polysaccharide antigen via MHC class II | 8.60e-04 | 3/13 |
| | 9 | **GO:1900117** | **regulation of execution phase of apoptosis** | **9.24e-09** | **5/12** |
| | | GO:2000272 | negative regulation of signaling receptor activity | 1.74e-08 | 5/12 |
| | | GO:0097194 | execution phase of apoptosis | 8.97e-07 | 5/12 |

Table S9: Gene Ontology enrichment analysis of cell-type-specific co-expressed gene modules identified with $\rho$-sctransform in B cells, CT4 T cells and CD8 T cells using PBMC samples from Wilk et al. (2020). Representative GO terms that capture cell-type-specific biological functions are highlighted in blue.

| Cell type | Module | GO term ID | GO term | Adjusted $p$-value | Gene ratio |
|---|---|---|---|---|---|
| B | 1 | GO:0045047 | protein targeting to ER | 8.04e-08 | 16/417 |
| | | GO:0072599 | establishment of protein localization to endoplasmic reticulum | 4.58e-07 | 16/417 |
| | | GO:0070085 | glycosylation | 7.08e-06 | 21/417 |
| | 2 | GO:0002181 | cytoplasmic translation | 8.44e-74 | 89/344 |
| | | GO:0006412 | translation | 4.81e-44 | 115/344 |
| | | GO:0043043 | peptide biosynthetic process | 4.81e-44 | 115/344 |
| | 3 | GO:0062023 | collagen-containing extracellular matrix | 2.48e-07 | 15/220 |
| | | GO:0030312 | external encapsulating structure | 4.57e-07 | 15/220 |
| | | GO:0031012 | extracellular matrix | 4.57e-07 | 15/220 |
| | 4 | **GO:0002399** | **MHC class II protein complex assembly** | **2.49e-14** | **13/195** |
| | | GO:0002503 | peptide antigen assembly with MHC class II protein complex | 2.49e-14 | 13/195 |
| | | GO:0019886 | antigen processing and presentation of exogenous peptide antigen via MHC class II | 2.66e-14 | 15/195 |
| | 5 | GO:0110053 | regulation of actin filament organization | 2.76e-05 | 17/142 |
| | | GO:0032970 | regulation of actin filament-based process | 2.76e-05 | 19/142 |
| | | GO:0032956 | regulation of actin cytoskeleton organization | 3.01e-05 | 18/142 |
| CD4T | 1 | GO:0030036 | actin cytoskeleton organization | 1.55e-05 | 52/474 |
| | | GO:0030029 | actin filament-based process | 1.55e-05 | 54/474 |
| | | GO:0030838 | positive regulation of actin filament polymerization | 1.55e-05 | 18/474 |
| | 2 | GO:0002181 | cytoplasmic translation | 4.94e-71 | 83/291 |
| | | GO:0006412 | translation | 1.36e-38 | 100/291 |
| | | GO:0043043 | peptide biosynthetic process | 1.60e-38 | 100/291 |
| | 3 | **GO:0002495** | **antigen processing and presentation of peptide antigen via MHC class II** | **1.24e-08** | **7/66** |
| | | GO:0019886 | antigen processing and presentation of exogenous peptide antigen via MHC class II | 1.24e-08 | 7/66 |
| | | GO:0002504 | antigen processing and presentation of peptide or polysaccharide antigen via MHC class II | 2.25e-08 | 7/66 |
| | 4 | GO:0008380 | RNA splicing | 2.09e-04 | 15/52 |
| | | GO:0016071 | mRNA metabolic process | 2.09e-04 | 18/52 |
| | | GO:0006397 | mRNA processing | 3.47e-04 | 15/52 |
| | 5 | **GO:0051607** | **defense response to virus** | **7.08e-12** | **17/55** |
| | | GO:0140546 | defense response to symbiont | 7.08e-12 | 17/55 |
| | | GO:0009615 | response to virus | 8.17e-11 | 18/55 |
| CD8T | 1 | GO:0051276 | chromosome organization | 2.14e-10 | 156/1071 |
| | | GO:0006260 | DNA replication | 8.55e-09 | 60/1071 |
| | | GO:0006281 | DNA repair | 1.60e-07 | 92/1071 |
| | 2 | GO:0046034 | ATP metabolic process | 1.36e-07 | 40/543 |
| | | GO:0097435 | supramolecular fiber organization | 2.03e-05 | 59/543 |
| | | GO:0007015 | actin filament organization | 2.03e-05 | 43/543 |
| | 3 | **GO:0019885** | **antigen processing and presentation of endogenous peptide antigen via MHC class I** | **9.80e-05** | **10/529** |
| | | GO:0002483 | antigen processing and presentation of endogenous peptide antigen | 5.93e-04 | 10/529 |
| | | GO:0019883 | antigen processing and presentation of endogenous antigen | 2.52e-03 | 10/529 |
| | 4 | GO:0002181 | cytoplasmic translation | 1.93e-78 | 86/271 |
| | | GO:0006412 | translation | 4.16e-41 | 99/271 |
| | | GO:0043043 | peptide biosynthetic process | 8.67e-41 | 99/271 |
| | 5 | **GO:0034976** | **response to endoplasmic reticulum stress** | **4.03e-05** | **10/35** |
| | | GO:0006457 | protein folding | 2.74e-04 | 8/35 |
| | | GO:0036503 | ERAD pathway | 2.22e-02 | 5/35 |

Table S10: Gene Ontology enrichment analysis of cell-type-specific co-expressed gene modules identified with $\rho$-sctransform in monocytes and natural killer (NK) cells using PBMC samples from Wilk et al. (2020). Representative GO terms that capture cell-type-specific biological functions are highlighted in blue.

| Cell type | Module | GO term ID | GO term | Adjusted $p$-value | Gene ratio |
|---|---|---|---|---|---|
| Monocyte | 1 | GO:0002181 | cytoplasmic translation | 5.34e-76 | 85/288 |
| | | GO:0006412 | translation | 2.21e-47 | 105/288 |
| | | GO:0043043 | peptide biosynthetic process | 5.63e-47 | 105/288 |
| | 2 | **GO:0002396** | **MHC protein complex assembly** | **4.22e-13** | **12/155** |
| | | GO:0002501 | peptide antigen assembly with MHC protein complex | 5.58e-12 | 11/155 |
| | | GO:0002399 | MHC class II protein complex assembly | 1.24e-11 | 10/155 |
| | 3 | **GO:0051607** | **defense response to virus** | **5.66e-24** | **33/106** |
| | | GO:0140546 | defense response to symbiont | 5.66e-24 | 33/106 |
| | | GO:0009615 | response to virus | 2.02e-20 | 34/106 |
| | 4 | GO:0014070 | response to organic cyclic compound | 5.05e-04 | 17/57 |
| | | **GO:0032623** | **interleukin-2 production** | **2.65e-03** | **6/57** |
| | | **GO:0032663** | **regulation of interleukin-2 production** | **2.65e-03** | **6/57** |
| | 5 | **GO:0006910** | **phagocytosis, recognition** | **2.26e-06** | **5/22** |
| | | GO:0006958 | complement activation, classical pathway | 2.62e-06 | 5/22 |
| | | GO:0002455 | humoral immune response mediated by circulating immunoglobulin | 6.65e-06 | 5/22 |
| NK | 1 | GO:0022610 | biological adhesion | 4.85e-04 | 64/432 |
| | | GO:0007155 | cell adhesion | 5.01e-04 | 63/432 |
| | | GO:0007186 | G protein-coupled receptor signaling pathway | 2.50e-03 | 28/432 |
| | 2 | **GO:0019885** | **antigen processing and presentation of endogenous peptide antigen via MHC class I** | **8.64e-06** | **9/282** |
| | | GO:0002483 | antigen processing and presentation of endogenous peptide antigen | 1.33e-05 | 9/282 |
| | | GO:0019883 | antigen processing and presentation of endogenous antigen | 5.52e-05 | 9/282 |
| | 3 | GO:0002181 | cytoplasmic translation | 2.42e-77 | 85/270 |
| | | GO:0006412 | translation | 2.46e-40 | 95/270 |
| | | GO:0043043 | peptide biosynthetic process | 5.33e-40 | 95/270 |
| | 4 | GO:0007015 | actin filament organization | 7.75e-09 | 23/120 |
| | | GO:0030036 | actin cytoskeleton organization | 7.75e-09 | 28/120 |
| | | GO:0030029 | actin filament-based process | 2.22e-08 | 28/120 |
| | 5 | **GO:0051607** | **defense response to virus** | **1.18e-07** | **12/39** |
| | | GO:0140546 | defense response to symbiont | 1.18e-07 | 12/39 |
| | | GO:0045071 | negative regulation of viral genome replication | 1.75e-06 | 7/39 |
| | 6 | GO:0006457 | protein folding | 2.42e-07 | 11/39 |
| | | **GO:0034976** | **response to endoplasmic reticulum stress** | **3.39e-07** | **12/39** |
| | | GO:0036503 | ERAD pathway | 3.79e-02 | 5/39 |
| | 7 | GO:0022616 | DNA strand elongation | 3.89e-04 | 4/36 |
| | | GO:0006270 | DNA replication initiation | 3.89e-04 | 4/36 |
| | | GO:0033260 | nuclear DNA replication | 3.89e-04 | 4/36 |

In Tables S11-S12, we evaluated the differential co-expression network estimated using CS-CORE and $\rho$-sctranform (see Section 4.6), respectively, using monocytes from control subjects and COVID patients in Wilk et al. (2020). We performed clustering analyses following WGCNA (Langfelder and Horvath, 2008) with the soft-thresholding power set to 1 to obtain differentially co-expressed gene modules. We then conducted GO enrichment analyses on the identified modules and the top (up to) three GO terms with the strongest enrichment signals are presented in Tables S11-S12 for modules with at least one significant GO term (BH-adjusted $p$-values $< 0.05$).

Table S11: Gene Ontology enrichment analysis of differentially co-expressed gene modules identified with CS-CORE in monocytes using PBMC samples from Wilk et al. (2020). Representative GO terms that capture cell-type-specific biological functions and/or cell-type-specific disease-related biological pathways are highlighted in blue.

| Cell type | Module | GO term ID | GO term | Adjusted $p$-value | Gene ratio |
|---|---|---|---|---|---|
| Monocyte | 1 | GO:0006935 | chemotaxis | 1.20e-02 | 19/101 |
| | | GO:0042330 | taxis | 1.20e-02 | 19/101 |
| | | **GO:0002532** | **production of molecular mediator involved in inflammatory response** | **2.03e-02** | **9/101** |
| | 2 | **GO:0051607** | **defense response to virus** | **4.69e-07** | **20/89** |
| | | GO:0140546 | defense response to symbiont | 4.69e-07 | 20/89 |
| | | GO:0009615 | response to virus | 8.09e-06 | 22/89 |
| | 3 | GO:0002181 | cytoplasmic translation | 1.38e-32 | 43/62 |
| | | GO:0006412 | translation | 1.36e-31 | 49/62 |
| | | GO:0043043 | peptide biosynthetic process | 1.36e-31 | 49/62 |
| | 4 | GO:0002181 | cytoplasmic translation | 7.44e-09 | 17/29 |
| | | GO:0006412 | translation | 3.77e-07 | 18/29 |
| | | GO:0043043 | peptide biosynthetic process | 3.77e-07 | 18/29 |
| | 5 | GO:0008092 | cytoskeletal protein binding | 2.44e-03 | 11/22 |
| | | GO:0051015 | actin filament binding | 2.00e-02 | 6/22 |
| | | GO:0003779 | actin binding | 3.31e-02 | 7/22 |
| | 6 | **GO:0080134** | **regulation of response to stress** | **4.98e-03** | **11/18** |
| | | GO:0043086 | negative regulation of catalytic activity | 4.98e-03 | 9/18 |
| | | GO:0009968 | negative regulation of signal transduction | 4.98e-03 | 9/18 |
| | 7 | GO:0101002 | ficolin-1-rich granule | 5.70e-03 | 7/17 |
| | | GO:1904813 | ficolin-1-rich granule lumen | 5.70e-03 | 6/17 |
| | | GO:0030141 | secretory granule | 5.70e-03 | 10/17 |
| | 8 | GO:0140534 | endoplasmic reticulum protein-containing complex | 2.29e-04 | 5/15 |
| | | GO:0071013 | catalytic step 2 spliceosome | 5.15e-03 | 4/15 |
| | | GO:0005681 | spliceosomal complex | 1.49e-02 | 4/15 |
| | 9 | GO:0003729 | mRNA binding | 9.73e-03 | 5/10 |

Table S12: Gene Ontology enrichment analysis of differentially co-expressed gene modules identified with $\rho$-sctransform in monocytes using PBMC samples from Wilk et al. (2020). Representative GO terms that capture cell-type-specific biological functions and/or cell-type-specific disease-related biological pathways are highlighted in blue.

| Cell type | Module | GO term ID | GO term | Adjusted $p$-value | Gene ratio |
|---|---|---|---|---|---|
| Monocyte | 1 | GO:0016192 | vesicle-mediated transport | 1.01e-03 | 101/428 |
| | | GO:0051179 | localization | 2.84e-03 | 227/428 |
| | | GO:0006810 | transport | 1.97e-02 | 178/428 |
| | 2 | GO:0050792 | regulation of viral process | 1.79e-03 | 19/202 |
| | | GO:0019079 | viral genome replication | 1.79e-03 | 17/202 |
| | | **GO:0051607** | **defense response to virus** | **1.79e-03** | **24/202** |
| | 3 | GO:0008022 | protein C-terminus binding | 6.61e-03 | 10/101 |
| | 4 | GO:0002181 | cytoplasmic translation | 1.38e-27 | 39/59 |
| | | GO:0006412 | translation | 1.40e-20 | 40/59 |
| | | GO:0043043 | peptide biosynthetic process | 1.40e-20 | 40/59 |
| | 5 | GO:0008092 | cytoskeletal protein binding | 3.56e-02 | 14/40 |
| | 6 | GO:0006457 | protein folding | 4.86e-03 | 6/24 |
| | | GO:0034976 | response to endoplasmic reticulum stress | 2.05e-02 | 6/24 |
| | | GO:0140534 | endoplasmic reticulum protein-containing complex | 4.69e-03 | 5/24 |
| | 7 | GO:0005525 | GTP binding | 5.72e-03 | 5/17 |
| | | GO:0019001 | guanyl nucleotide binding | 5.72e-03 | 5/17 |
| | | GO:0032561 | guanyl ribonucleotide binding | 5.72e-03 | 5/17 |
| | 8 | GO:0043202 | lysosomal lumen | 2.28e-02 | 4/18 |

Finally, for the differentially co-expressed gene module presented in Figure 5, we further used hierarchical clustering to generate three sub-modules and annotated the functions of each sub-module with GO and Reactome enrichment analyses (see Sections S3.1), with background set to all genes in the databases to interpret the gene sets. Representative GO terms and Reactome pathways are reported for each sub-module.

Table S13: Gene Ontology and Reactome enrichment analysis for the three sub-modules in Figure 5.

| Module | Pathway | ID | Description | Adjusted $p$-value | Gene ratio |
|---|---|---|---|---|---|
| 1. Toll-like receptor signalling | GO | GO:0050665 | hydrogen peroxide biosynthetic process | 2.13e-03 | 2/5 |
| | | GO:1903428 | positive regulation of reactive oxygen species biosynthetic process | 2.13e-03 | 2/5 |
| | | **GO:0034121** | **regulation of toll-like receptor signaling pathway** | **5.55e-03** | **2/5** |
| | Reactome | R-HSA-3299685 | Detoxification of Reactive Oxygen Species | 1.14e-02 | 2/6 |
| 2. Interferon signalling | GO | GO:0009615 | response to virus | 7.22e-23 | 17/21 |
| | | GO:0051607 | defense response to virus | 3.14e-21 | 15/21 |
| | | GO:0140546 | defense response to symbiont | 3.14e-21 | 15/21 |
| | Reactome | R-HSA-909733 | Interferon alpha/beta signaling | 2.03e-14 | 9/18 |
| | | **R-HSA-913531** | **Interferon Signaling** | **3.27e-14** | **11/18** |
| | | R-HSA-1169410 | Antiviral mechanism by IFN-stimulated genes | 2.96e-08 | 6/18 |
| 3. Antigen Presentation | GO | **GO:0019885** | **antigen processing and presentation of endogenous peptide antigen via MHC class I** | **4.42e-04** | **3/18** |
| | | GO:0002483 | antigen processing and presentation of endogenous peptide antigen | 4.42e-04 | 3/18 |
| | | GO:0019883 | antigen processing and presentation of endogenous antigen | 7.87e-04 | 3/18 |
| | Reactome | R-HSA-1236977 | Endosomal/Vacuolar pathway | 3.29e-04 | 3/19 |
| | | R-HSA-983170 | Antigen Presentation: Folding, assembly and peptide loading of class I MHC | 2.26e-03 | 3/19 |
| | | **R-HSA-9694516** | **SARS-CoV-2 Infection** | **1.59e-02** | **3/19** |

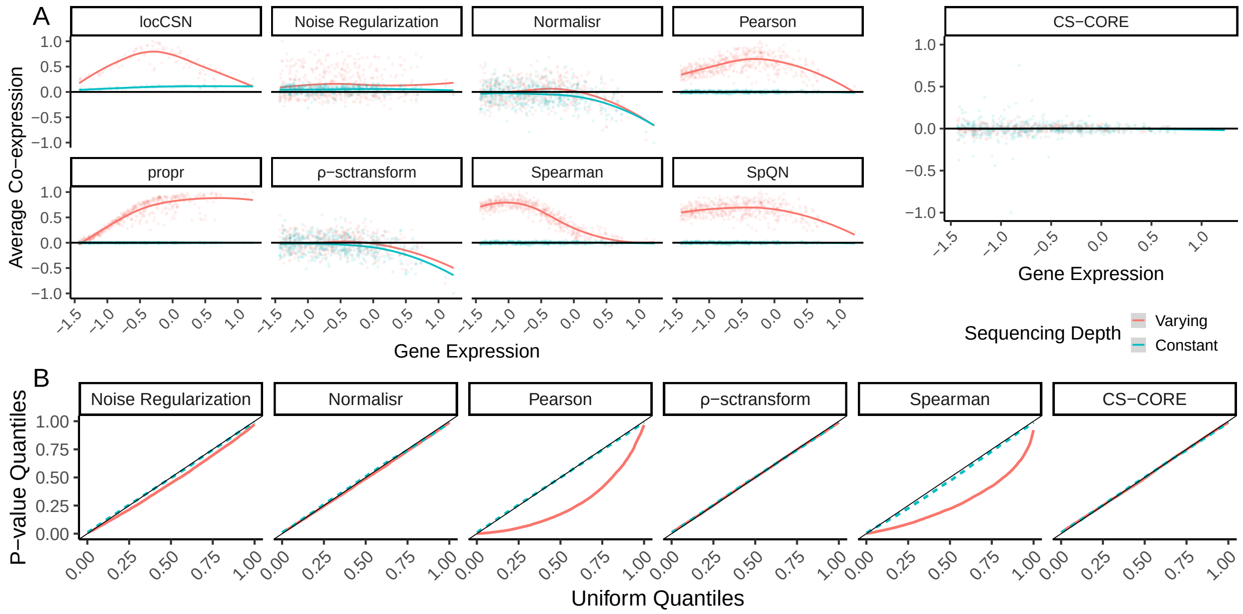


Figure S2: Validation of CS-CORE using simulated scRNA-seq data with independent gene expressions. We selected genes and cells as in Figure 1 and simulated data as described in Section 4.5 with a diagonal correlation matrix and marginal negative binomial distributions. Results from simulated data with varying and constant sequencing depths are colored with light red and blue, respectively. (A) Scatter plots with fitted curves showing expression (x-axis) and average co-expression (y-axis) of each gene with co-expression estimated using locCSN, Noise Regularization, Normalisr, Pearson correlation, propr, $\rho$-sctransform, Spearman correlation, SpQN and CS-CORE. Average co-expressions are re-scaled by the maximum value to aid comparison. (B) Q-Q plots comparing $p$-values for testing co-expressions of gene pairs against Uniform(0,1) using six methods with statistical tests, including Noise Regularization, Normalisr, Pearson correlation, $\rho$-sctransform, Spearman correlation and CS-CORE.

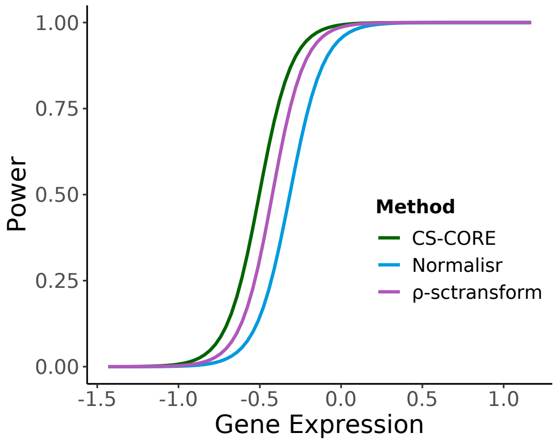


Figure S3: Statistical power in detecting co-expression at different gene expression levels. The three methods being compared are the ones that have appropriate type-I error controls in Figure 1B, including CS-CORE, Normalisr and $\rho$-sctransform. Data were simulated under the same setting as in Figure 2A.

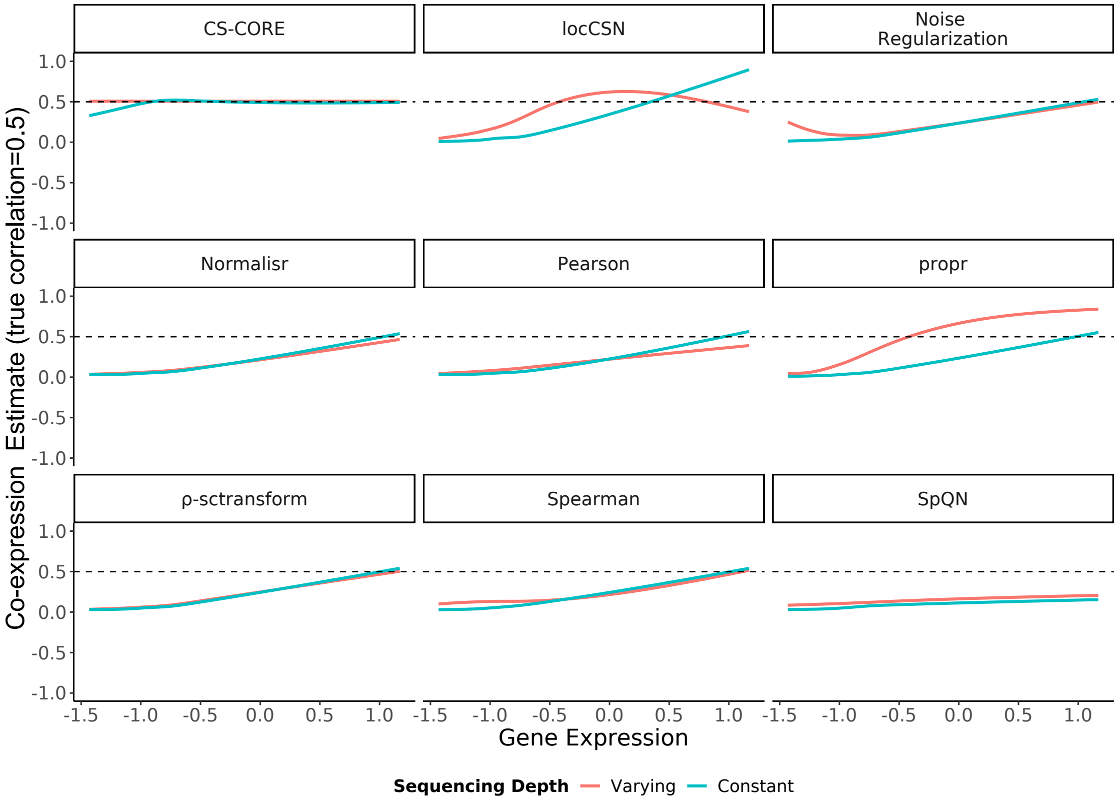


Figure S4: Validation of CS-CORE co-expression estimates using simulated data with constant and varying sequencing depths, compared to locSCN, Noise Regularization, Normalisr, Pearson and Spearman correlations, propr, $\rho$-sctransform and SpQN. Curve-fitted co-expression estimates are plotted against geometric mean expression levels on gene pairs simulated with a true correlation of 0.5 (5,000 genes and 1,000 cells). Data were simulated as described in Section 4.5.

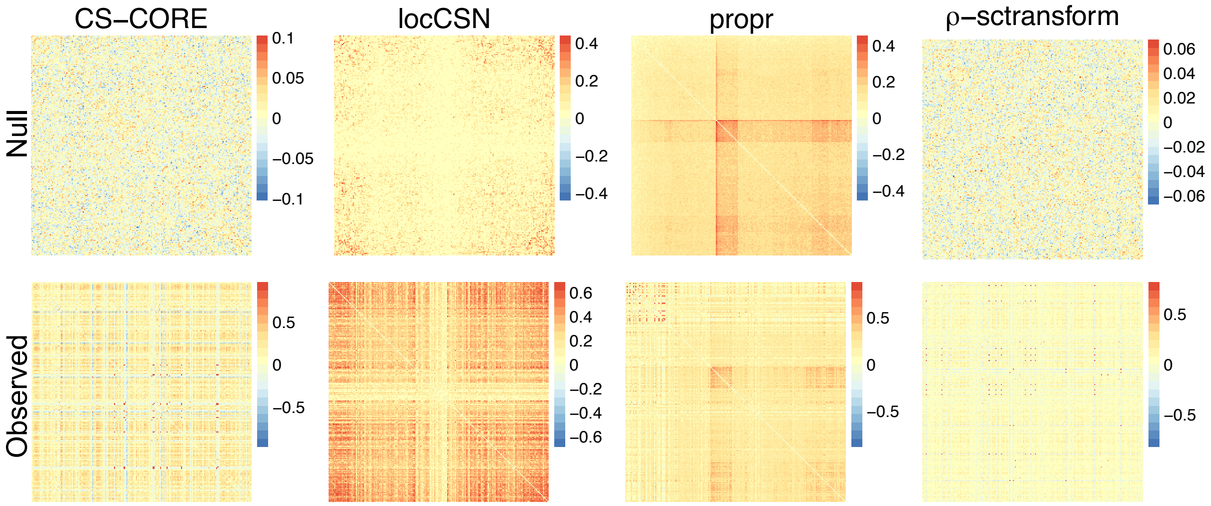


Figure S5: Clustering structures on permuted null data and observed data for CS-CORE, locCSN, propr and $\rho$-sctransform. We estimated co-expression networks for the top 200 highly expressed genes using both permuted null data and observed data on excitatory neurons of control subjects from Lau et al. (2020), where the permuted null data were generated as in Figure 1 and all gene expressions should be independent in the permuted null data. For estimates of the same method, the genes are ordered by hierarchical clustering of the estimates based on null data.

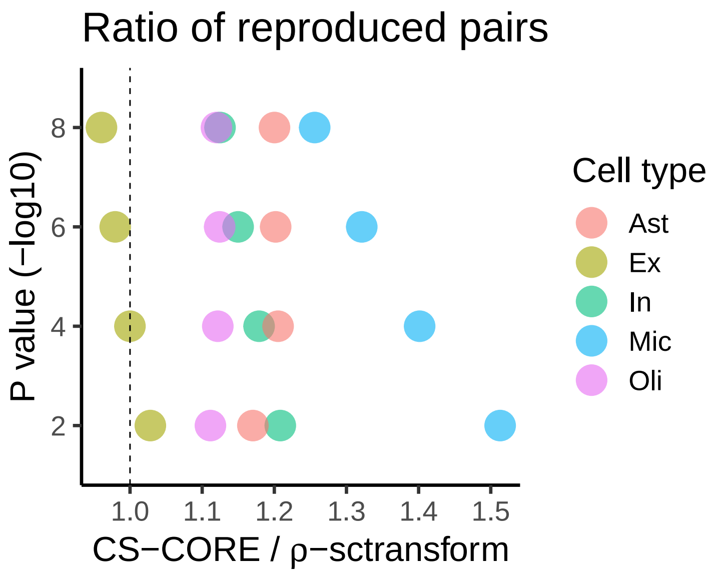


Figure S6: Ratio of the numbers of gene pairs that were identified as significant in both Lau et al. (2020) and Morabito et al. (2021) at specified $p$-value cutoffs between CS-CORE and $\rho$-sctransform. We used the cells in five major brain cell types from control subjects from Lau et al. (2020) and Morabito et al. (2021) respectively to estimate cell-type-specific co-expression networks.

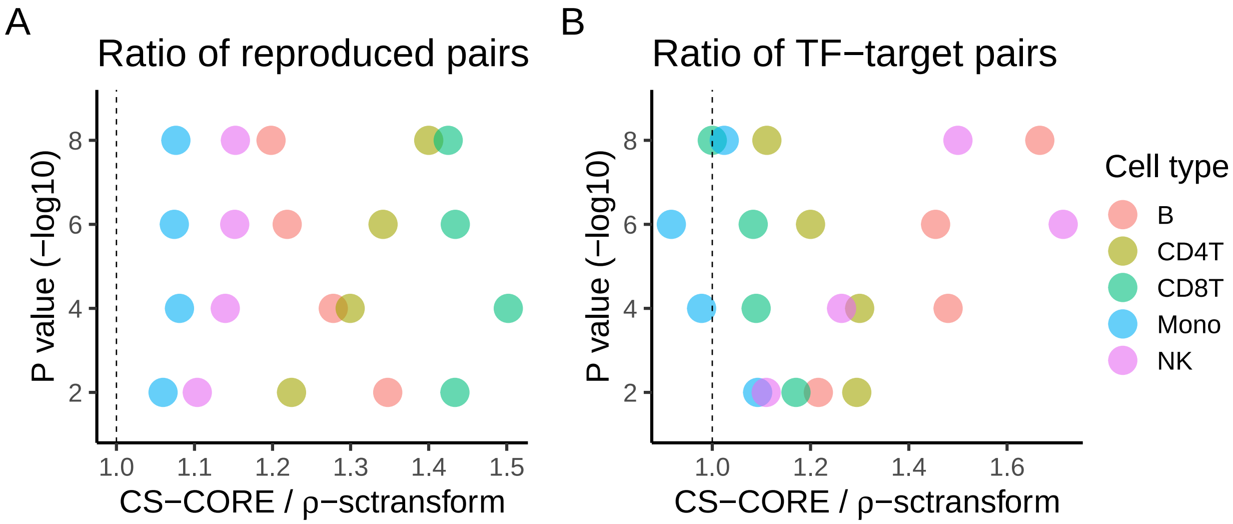


Figure S7: Co-expression analysis using PBMC samples in Wilk et al. (2020). We used the cells in five major immune cell types from control subjects from Wilk et al. (2020) to estimate cell-type-specific co-expression networks. **A.** Ratio of the numbers of gene pairs that were identified as significant in both Wilk et al. (2020) and Unterman et al. (2022) at specified $p$-value cutoffs between CS-CORE and $\rho$-sctransform. **B.** Ratio of the numbers of gene pairs that were identified as significant and overlapped with known TF-target gene pairs in the TRRUST database (Han et al., 2018) between CS-CORE and $\rho$-sctransform.

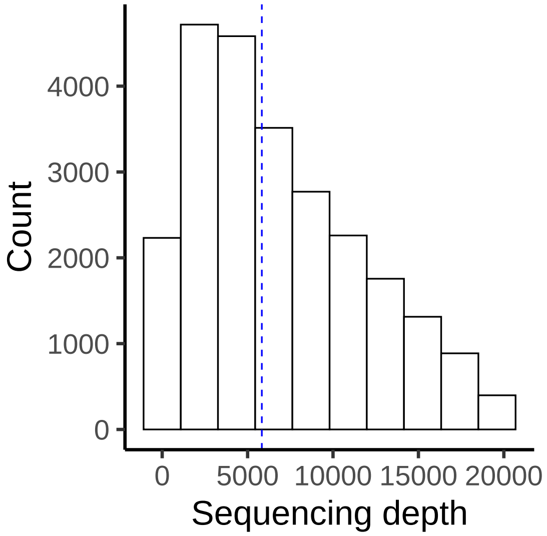


Figure S8: Histogram of sequencing depths (sum of UMI counts in a cell) of excitatory neurons from control subjects from Lau et al. (2020). The median 5,833 is marked with a dashed blue line.

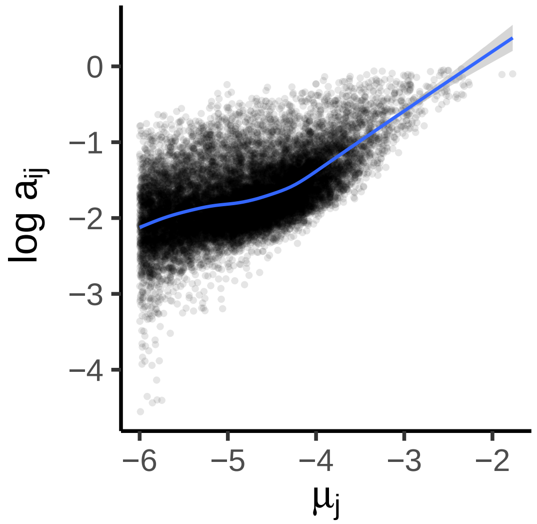


Figure S9: Log attenuation factor $\log a_{ij}$ across expression levels $\mu_j$ in cells with sequencing depth $s_i = 2{,}000$. Marginal estimates of $\mu_j$ and $\mathrm{CV}_j$ were obtained as described in Section S2 and $a_j$ was calculated as $\sqrt{(s_i\mathrm{CV}_j^2)/(1/\mu_j + s_i\mathrm{CV}_j^2)}$.
